# Trainable computation in molecular networks

**DOI:** 10.64898/2025.12.28.696421

**Authors:** Kristina Trifonova, Martin J. Falk, Mason Rouches, Suriyanarayanan Vaikuntanathan, Michael Elowitz, Arvind Murugan

## Abstract

Reports of learning in single cells without genetic change span decades yet remain controversial, in part because there is no accepted general molecular mechanism for training comparable to gradient-based training or Hebbian learning in neural circuits. Here we identify a minimal set of ingredients sufficient to realize non-genetic learning, drawing inspiration from Boltzmann neural networks. First, dense reversible interaction networks provide an expressive substrate in which modulating the concentrations of a small set of mediator species can reprogram function without altering the underlying interaction parameters. Second, a simple rate-sensitive autoregulatory scheme that adjusts these mediator levels provides a local Hebbian-like training rule that can train the same network for diverse tasks, including Pavlovian conditioning, supervised classification, and generative tuning of bet-hedging ratios to match environmental statistics. We show that this autoregulatory training rule is model free and applies to reversible multimerization networks of arbitrary complexity, so training can compensate for unknown or unmodeled interactions present in vivo. These results suggest design principles for trainable synthetic cellular circuits and indicate how molecular systems could learn statistical features of their environments through experience.

---

Natural selection is often invoked to explain how cells come to embody the statistics of their past environments. In this view, regulatory programs encode those statistics, shaping cellular responses based on how frequently different conditions occurred in the past and not just on their immediate presence. These evolved programs can be remarkably sophisticated: microbes can anticipate nutrient depletion [1–4], populations hedge their bets by adopting diverse phenotypes at characteristic frequencies [5–9], and stress response pathways activate in anticipation of correlated environmental changes [10–12]. Here, memory of environmental statistics is genetically hardwired over evolutionary timescales through mutation and selection.

Yet many observations hint that cells can learn environmental statistics on much shorter timescales, via parameter remodeling within a lineage rather than differential survival of variants. These observations [8, 13–26] include habituation, priming, Pavlovian associative conditioning, anticipatory regulation, and learned bet hedging in cells ranging from bacteria to protists, yeast, and immune and cancer cells. While intriguing, many of these observations remain controversial, especially since the mechanisms underlying these phenomena are not fully clear.

Here, we define learning as experience-dependent reprogramming of regulatory programs: repeated exposure to a structured environment adjusts internal parameters so that future responses reflect correlations in past stimuli rather than only their immediate presence [27–29]; see Fig.1a. Biologically, learning within a lifetime would allow cells and organisms to adapt to evolutionarily novel or fluctuating environments on short timescales and without the cost of deleterious mutations. In a synthetic biology context, it would enable the engineering of robust synthetic circuits that are trained *in situ* to perform useful behaviors, alleviating the need for mechanistic models or repeated rounds of fine-tuning.

**FIG. 1.**
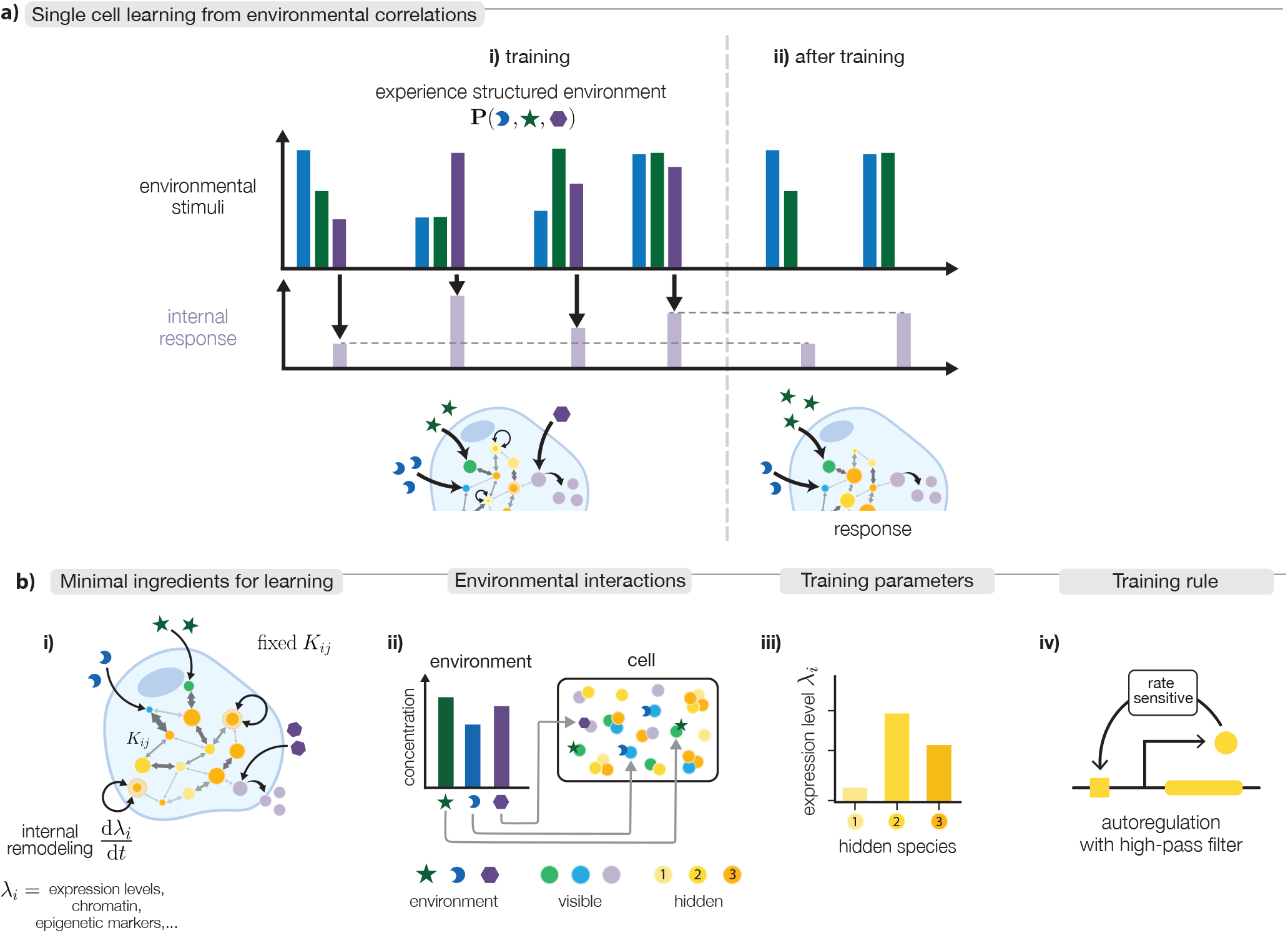
Trainable molecular networks enable single cells to learn environmental correlations. (a) Operational definition of learning. Bars show the levels of several environmental inputs (blue, green, purple) that fluctuate jointly according to a structured distribution *P* (☽, ★, ⬡). We focus on an internal response (light purple) induced directly by the purple stimulus (hexagon). During exposure to a structured environment (i), a cell repeatedly experiences correlated combinations of stimuli that drives changes in its internal molecular parameters. After training (ii), blue and green inputs by themselves elicit an internal response that approximates the response that would occur if the purple cue were present; such a molecular network is said to have learned the statistical structure of environmental stimuli. (b) Minimal molecular ingredients for learning: (i) We consider a dense reversible interaction network between many molecular species with fixed microscopic interaction parameters *K*_*ij*_. Training acts by slowly tuning other internal parameters *λ*_*i*_ according to a training rule while leaving *K*_*ij*_ unchanged. (ii) Environmental molecules impact a subset of internal “visible” species, shifting the equilibrium of the internal network. Other internal components, “hidden” species, do not couple directly to the environment but influence cellular response. (iii) In our study, concentrations of hidden species play the role of training parameters *λ*_*i*_. (iv) Training rule: Hidden species levels are trained by a rate-sensitive autoregulatory rule: rapid changes in the concentration of a monomer drives production or degradation of that species, whereas steady monomer levels do not (i.e., autoregulation with high-pass filtering).

At present, no general mechanism is known by which molecular systems can learn from experience across diverse molecular architectures and learning behaviors. In comparison to learning in neural systems, learning in non-neural cells has seemed mechanistically implausible because of two fundamental obstacles. First, the relevant internal parameters, analogs of synaptic weights in neural networks, must be tunable without mutations, which rules out direct molecular affinities as potential training parameters. Second, a molecular ‘training rule’ must locally adjust these parameters to improve performance, without access to global computations such as back-propagation that enable learning in artificial neural networks.

Here, we show that both obstacles can be overcome in reversible molecular networks in which each species interacts with many others. In such densely interacting networks, the total concentrations of a small set of ‘hidden’ mediator species can serve as trainable variables that are readily modulated by regulation without requiring mutations [28, 30–33]. Moreover, adapting training ideas from Boltzmann machines [34, 35] to the molecular domain, we show that simple rate-sensitive autoregulation of these hidden species yields a general training rule that tunes their concentrations toward values consistent with environmental correlations experienced during training; see Fig.1b. Across different environmental contexts, this rate-sensitive autoregulatory rule enables a wide class of reversible molecular networks to undergo Pavlovian conditioning, learn quantitative input–output relationships, perform supervised classification, and learn generative behaviors relevant for bet hedging. We argue that the required molecular ingredients, dense interaction networks, autoregulation, and rate sensitivity, are widespread in biology. Our findings therefore suggest a broadly applicable, model-free mechanism by which molecular systems can reprogram their function on physiological timescales without requiring genetic change.

## I. TRAINING FOR PAVLOVIAN CONDITIONING

We begin with the classic problem of Pavlovian conditioning [14, 36], in which a neutral stimulus becomes predictive of a biologically potent one. Pavlovian conditioning in non-neural contexts has been claimed since Gelber’s early work [37] on protists, with subsequent observations in ciliates and amoebae, slime molds and plants, and, more recently, in macrophages and in cancer drug resistance [17, 38–47]. However, alternative explanations and the lack of clear molecular mechanisms render many of these claims controversial [13, 22]. Prior work has explored diverse molecular routes to learning and memory, from evolutionary and phenomenological proposals to task-specific circuits [1, 2, 28, 29, 48–59]. Here we ask whether a minimal, physically grounded mechanism for such associative learning can be built from reversible molecular binding.

In our molecular Pavlovian setup, an environmental signal *E*_*B*_ acts as the *neutral stimulus* (“bell”) and another *E*_*F*_ as the *potent stimulus* (“food”) (Fig. 2). These signals modulate internal species *V*_*B*_ and *V*_*F*_, whose reversible associations with other species, for example through dimerization [31, 60] or higher-order multimerization [32, 33, 35, 61], provide the signal-processing substrate for the learned response. For concreteness, we take 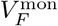 as the functional response, although variants where 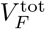 or 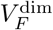 serve as response behave equivalently (SI Sec. IV). Initially, *E*_*B*_ alone produces no consistent *V*_*F*_ response; training modifies the molecular network so that repeated co-presentation of *E*_*B*_ and *E*_*F*_ makes *E*_*B*_ alone elicit the response previously induced by *E*_*F*_.

**FIG. 2.**
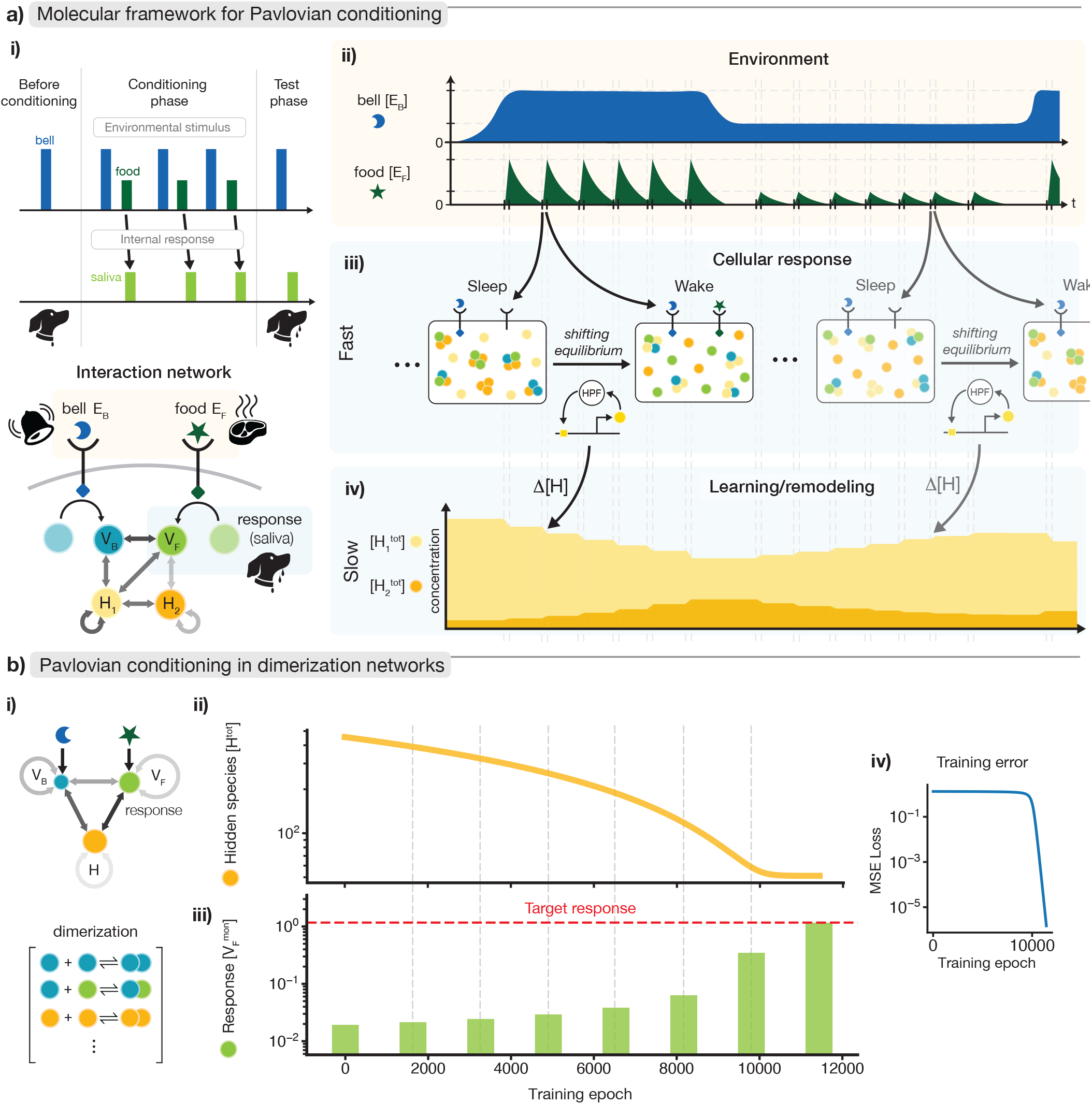
Pavlovian conditioning through autoregulation of hidden species in a structured environment. **a**. (i) Schematic of molecular Pavlovian conditioning. (ii) The neutral and potent stimuli (*bell* and *food*) are environmental molecules *E*_*B*_ and *E*_*F*_ that regulate internal species *V*_*B*_ and *V*_*F*_ within a network containing hidden species (*H*_1_, *H*_2_). (iii) Training uses correlated inputs: in the example shown, *E*_*F*_ spikes to an *E*_*B*_-dependent level, rapidly driving the network from a baseline ‘sleep’ state to an *E*_*F*_-driven ‘wake’ state before slowly returning after *E*_*F*_ is removed. (iv) Hidden-species totals 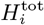 are the trainable parameters, updated by rate-sensitive autoregulation of each species’ own monomer level 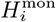. Training drives these totals toward values that reduce the sleep-wake mismatch, so that *E*_*B*_ alone reproduces the response previously induced by the correlated *E*_*F*_ spike. **b**. (i) Simulation results of Pavlovian conditioning in a three-species dimerization network (*V*_1_, *V*_2_, *H*) coupled to two stimuli *E*_*F*_, *E*_*B*_. Grayscale values of network edges represent strengths of binding constants *K*_*ij*_ which stay fixed during training. (ii) Accumulation of the hidden species *H*^tot^ in response to spikes of *E*_*F*_ that occur only when *E*_*B*_ is high during training. (iii) Testing during training: output response 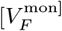 to periodic test pulses of the neutral-stimulus molecule *E*_*B*_ presented without *E*_*F*_. As learning proceeds, these test responses increase toward the desired level. (iv) Mean-squared error (MSE) between actual and target output 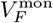 in response to *E*_*B*_ decreases over training epochs, demonstrating Pavlovian conditioning.

We assume that each environmental signal *E*_*X*_ (e.g., *E*_*B*_ or *E*_*F*_) acts through a receptor-mediated pathway that directly regulates a corresponding monomer concentration 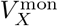. Common biological mechanisms include *E*_*X*_-activated receptors that catalytically promote reversible exchange of 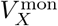 with a buffered pool, for example through sequestration into an inactive state or trafficking into another compartment. When *E*_*X*_ is high, this Sflux dominates the slower constitutive processes that would otherwise determine the ambient level of *V*_*X*_ and sets 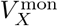 to a *E*_*X*_-dependent value. When *E*_*X*_ is low or absent, the receptor-mediated flux weakens or vanishes, and intrinsic homeostatic dynamics restore the broader *V*_*X*_ pool, in the simplest case 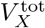, toward a fixed baseline. Mathematical models of this environmental coupling and alternative compatible couplings are discussed in the SI.

We now distinguish between species directly controlled by environmental signals and those affected only through the molecular network. The former, *V*_*B*_ and *V*_*F*_, play the role of *visible* species in the language of Boltzmann machines [34, 35, 62], while the latter, *H*_*i*_, are *hidden* species (Fig. 2a). These species can reversibly form dimers, trimers, and higher-order complexes through fixed affinities *K*_*ij*…_, providing the equilibrium binding network that processes environmental inputs. Rather than changing affinities during learning, our model trains hidden-species totals 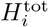, because cellular concentrations can change dynamically through synthesis and degradation whereas molecular affinities are fixed on the timescale of an individual’s experience. The training rule below uses only local changes in each hidden species’ monomer level and is compatible with many network topologies (SI Sec. II).

To induce training, we use a structured environment in which *E*_*F*_ appears in spikes whose height depends on whether *E*_*B*_ is high or low (Fig. 2b). We refer to periods with no *E*_*F*_ as “sleep” and periods with *E*_*F*_ present as “wake” [34, 35, 62]. A key requirement for learning is temporal asymmetry [63]: *E*_*F*_ onset rapidly drives the system from sleep to wake on a timescale *τ*_*f*_, whereas return to sleep occurs slowly over *τ*_*s*_. Such asymmetry can arise from fast ligand appearance followed by slow dilution or degradation, or from slow constitutive restoration of 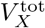 after *E*_*F*_-driven receptor activity ends.

The trainable variables 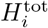 are updated by a local autoregulatory rule derived by adapting Boltzmann-learning ideas to molecular systems [34, 35, 62, 63] (SI Secs. I, V). Each hidden species responds only to rapid changes in its own monomer level 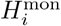 : slow or steady levels are ignored, while abrupt sleep-to-wake changes drive

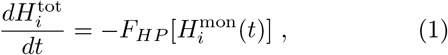

where *F*_*HP*_ is an effective high-pass filter with timescale *τ*_*HP*_. Because wake-to-sleep relaxation is slow compared with *τ*_*HP*_, the rule primarily samples the fast sleep-to-wake transition. Familiar adaptive biochemical motifs and biophysical mechanisms can result in such high-pass filtering (SI Sec. V).

We demonstrate this mechanism in a minimal three-species dimerization network (Fig. 2b) comprising visible species *V*_*B*_, *V*_*F*_, and one hidden species *H* coupled by fixed affinities *K*_*ij*_. Suppose the system starts with high [*H*^tot^]. In the sleep state with *E*_*B*_ present and *E*_*F*_ = 0, the output 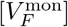 is too low because *H* binds *V*_*F*_ and depletes *V*_*F*_ monomers. During wake, the *E*_*F*_-driven pathway imposes the high-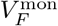 state; because *V*_*F*_ binds *H*, this perturbation also shifts *H* into *V*_*F*_-*H* dimers and lowers [*H*^mon^]. This decrease acts as an error signal: the autoregulatory rule decreases *H*^tot^, reducing *V*_*F*_ sequestration in subsequent sleep states.

Through repeated sleep-wake cycles, [*H*^tot^] decreases until *H*^mon^ become nearly invariant between the two states. The neutral stimulus *E*_*B*_ alone then elicits the response previously requiring *E*_*F*_ : the deviation of 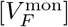 from its target falls to near zero over training epochs (Fig. 2b, right), while [*H*^tot^] decreases by almost an order of magnitude, encoding the learned association in steady state.

This adaptive feedback mechanism is broadly applicable: any reversible molecular network whose equilibrium distribution shifts under environmental inputs can, in principle, be trained by the same local autoregulatory rule. The expressive capacity of such systems, set by the fixed affinity structure *K*_*ij*_, is conceptually distinct from the question of how such systems learn *in situ*. Our framework above requires *τ*_*eq*_ *< τ*_*f*_ *< τ*_*s*_, with the filtering timescale *τ*_*HP*_ between the fast and slow environmental dynamics (SI Sec. V for a detailed discussion).

## II. SUPERVISED TRAINING FOR INPUT-OUTPUT BEHAVIORS

The same framework extends from Pavlovian associations to quantitative input-output maps between environmental inputs *E*_*B*_ and molecular responses 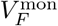. The autoregulatory rule is unchanged; only the training environment needs to be changed. In Pavlovian conditioning, the bell *E*_*B*_ is trained to induce whatever response 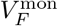 is naturally associated with a level of *E*_*F*_. In supervised training, the environment instead supplies a teaching signal: for each input level *E*_*B*_, the amplitude of the accompanying *E*_*F*_ pulse is chosen so that 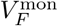 is driven to a desired target value. Such control could result from closed-loop feedback that adjusts *E*_*F*_ until the target 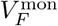 is reached.

With this supervised protocol, the same molecular networks can learn quantitative input *E*_*B*_-output 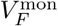 relations. For example, when low and high levels of *E*_*B*_ are paired with high and small *E*_*F*_ pulses, respectively, the three-species dimerization circuit learns a step-down relationship between *E*_*B*_ and 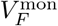. An eight-species network can learn a nonmonotonic low-high-low mapping when trained with correspondingly nonmonotonic *E*_*F*_ amplitudes, and can also be trained for a two-input function when exposed to two input signals, *E*_1_ and *E*_2_, together with the teaching signal *E*_*F*_ (Fig. 3c). Implementation details are provided in SI Sec. VI.

**FIG. 3.**
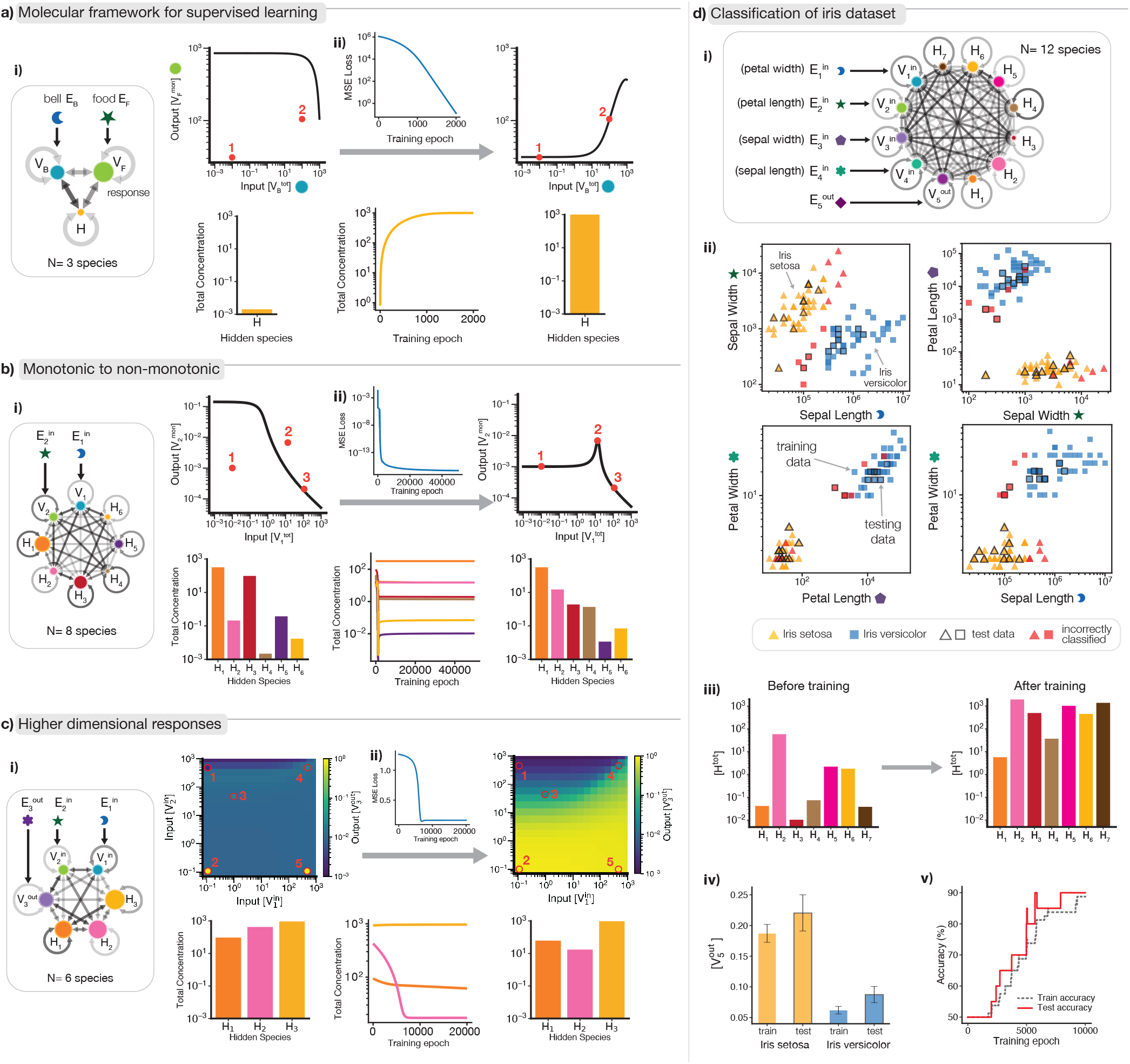
Supervised molecular learning of quantitative and multidimensional input–output behaviors. **a**. (i) Schematic of the training environment for supervised learning. Environmental inputs *E*_*B*_ (“bell”) and *E*_*F*_ (“food”) vary in correlated patterns such that high *E*_*B*_ coincides with high amplitude *E*_*F*_ spikes (training point 1) and vice versa (training point 2). (ii) A three-species network (*V*_*B*_, *V*_*F*_, *H*) initially shows an incorrect input–output relation between 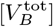 and 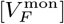. During training, hidden-species levels adjust themselves through the autoregulation training rule (Eq.1), reducing deviation from the target mapping. After training, the network reproduces the environmental correlations: high *E*_*B*_ yields high *V*_*F*_, and low *E*_*B*_ yields low *V*_*F*_. **b**. Extension to more complex mappings. (i) An eight-species network learns a non-monotonic response of output 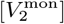 to input 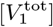. (ii) Internal concentrations evolve and the mean-squared error (MSE) from the target output decreases over training. **c**. The same rule generalizes to multidimensional inputs: (i) an eight-species network learns a 2D input–output function mapping 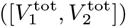 to 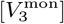. (ii) Training data shown in red circles. **d**. Classification of the Iris dataset using a twelve-species network. (i) Four input features (sepal and petal length and width) are encoded in stimuli (*E*_1_, *E*_2_, *E*_3_, *E*_4_) which induce concentrations of visible species 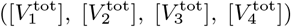, and the output molecule 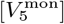 represents flower identity (low for *I. setosa*, high for *I. versicolor*). After training, (ii) the network reorganizes hidden species concentrations to (iii) separate species (triangle is *setosa*, square is *versicolor*, and red points are classified as the incorrect species). Training results in high 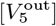 for *setosa* and low 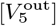 for *versicolor*, classifying points with (iv) ~89% accuracy on the training set and ~90% accuracy on the test set and demonstrating that the same molecular training rule can generalize to high-dimensional classification tasks.

To test whether the same rule scales to higher-dimensional inputs, we applied it to the standard four-feature Iris classification task, treating the four feature values as environmental signals *E*_1_, …, *E*_4_. After training on 80 examples from two classes, the output species classified *setosa* and *versicolor* with ~89% training accuracy and ~90% test accuracy (Fig. 3d). Thus, the same local molecular rule supports generalization beyond simple associative tasks.

## III. TRAINING FOR A DISTRIBUTION OF PHENOTYPES

The same autoregulatory training rule also extends to circuits whose behavior is stochastic and distributional rather than a deterministic input-output map. Many biological systems exhibit stochastic phenotypic variability [5], in which genetically identical cells occupy distinct states at characteristic frequencies and switch between them over time. Such controlled variability supports bet hedging across diverse settings, including bacterial persistence [64], yeast stress-activation heterogeneity [7], developmental fate diversification [65, 66], phage lysis– lysogeny [67], and bacterial competence [68, 69].

The occupancy fractions of different phenotypes are generally assumed to be genetically hardwired by evolution to match long-term environmental frequencies [6]; these fractions have also been hardwired in synthetic circuits [70, 71]. However, several studies [7–9, 18, 23, 24, 72] suggest that such fractions can instead be learned from experience, but a general molecular mechanism for tuning phenotypic distributions remains unknown. Here we show that our rate-sensitive autoregulatory rule provides such a mechanism.

Bistable molecular circuits provide a natural substrate for generating distributions of phenotypes, since intrinsic noise drives transitions between two attractor states. In many natural and synthetic examples (Fig. 4), the occupancy of these states can be tuned by adjusting the level of a specific molecular species. This suggests a simple principle: regulating the concentration of such a species with a training rule can train the stationary distribution.

**FIG. 4.**
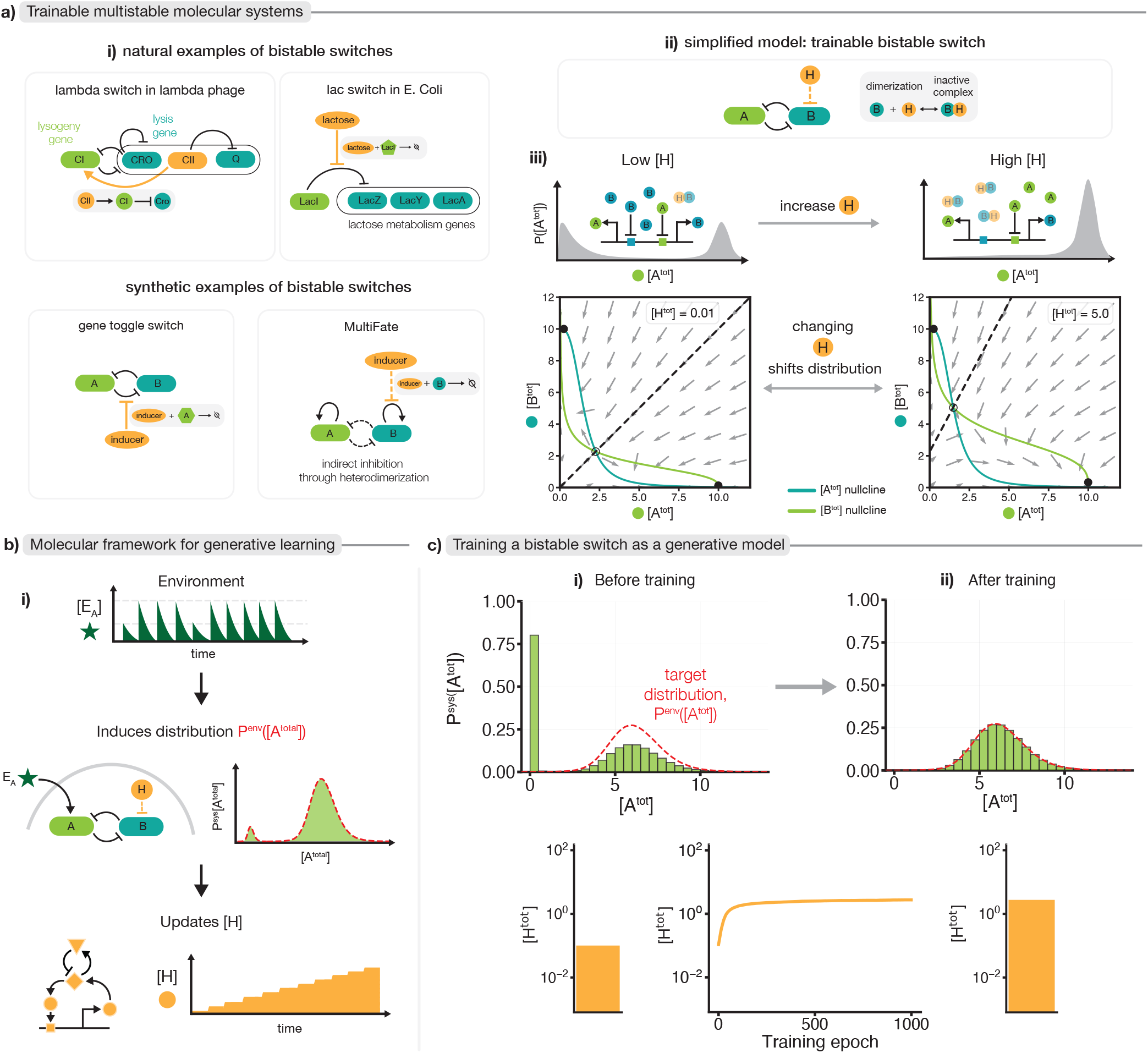
Training phenotype occupancy of a noisy bistable switch to match environmental statistics. **a**. (i) Examples of natural [73, 74] and synthetic [75, 76] bistable molecular switches in which the concentration of one molecule (orange) tunes the relative stability of two attractor states. (ii) A trainable bistable model composed of two mutually inhibiting species (*A, B*); a hidden species *H* sequesters *B* into an inactive complex. The two stable states correspond to high-*A*/low-*B* and low-*A*/high-*B* configurations. (iii) Probability distributions *P* ^sys^([*A*^tot^]) in the presence of intrinsic noise and phase-plane plots for low and high [*H*^tot^]; changing [*H*^tot^] shifts the relative stability and thus occupancy of these states. **b**. (i) During training, an environmental signal *E*_*A*_ drives the system into the high and low *A* states at frequencies *P* ^env^([*A*]^tot^) (dashed red line; schematic) defined by the environment. The autoregulatory training rule adjusts the hidden species [*H*^tot^] (orange; schematic) based on changes in [*H*^mon^]. **c**. (i) Environmental distribution *P* ^env^([*A*^tot^]) (red) and the noisy bistable switch’s distribution *P* ^sys^([*A*^tot^]) (green), with initial untrained value of *H*^tot^. Training of *H* was carried out as illustrated in (b). (ii) After training, the system’s state-occupancy distribution *P* ^sys^([*A*^tot^]) matches the environmental distribution *P* ^env^([*A*^tot^]); levels of hidden species *H* change by ~ 10*×* during training.

We test this idea using a minimal bistable network composed of two mutually inhibitory species (*A, B*), whose relative balance is controlled by a hidden species *H* that sequesters *B* into an inactive complex (Fig. 4a). Increasing [*H*^tot^] stabilizes the high-*A*/low-*B* state, while decreasing it shifts the balance towards the high-*B* state. Although this probabilistic behavior differs qualitatively from associative or supervised learning, SI Sec. II shows that the Boltzmann-learning framework predicts that autoregulation based on [*H*^mon^] will tune the stationary occupancy of the two attractors to match experienced environmental frequencies.

In simulations, we initialize the switch with an occupancy bias (i.e., level of [*H*^tot^]) that differs from the environment (e.g., 50*/*50 vs. 10*/*90 for the low- and high-*A* states; SI Sec. II). During training, an environmental signal *E*_*A*_ drives the system into the two configurations with frequencies *P* ^env^([*A*]^tot^) that reflect environmental statistics, inducing fluctuations in *H*^mon^ that in turn autoregulate *H*^tot^. After repeated training cycles, [*H*^tot^] converges to a value that spontaneously reproduces the environmental occupancy fractions experienced in the past, even in the absence of any environmental signal *E*_*A*_ (Fig. 4b). In a cellular population, this corresponds to a learned allocation strategy in which the fraction of cells adopting each phenotype reflects the statistics of previously encountered environments.

## IV. THE TRAINING FRAMEWORK IS MODEL-FREE

Thus far, we have demonstrated our auto-regulatory training framework on dimerization networks and bistable switches. Since this rule was inspired by Boltzmann machine training, a natural question arises: can we derive training rules that work for more complex molecular systems? Many biological networks involve reversible associations beyond simple dimerization, including trimerization, higher-order complexes, or even liquid condensates. Would each such molecular system require a distinct training rule, or could the same training mechanism apply broadly?

In this section, we show that the training framework is model-free: the training rule has the same functional form across reversible dilute association networks and relies only on measured concentration signals (and their transients). As a result, the training rule can be applied without knowing the physics of the underlying network (e.g., dimerization vs multimerization), interaction parameters, and remains effective under unmodeled interactions and cell-to-cell variability. In SI Sec. II, we derive the training rule for molecular networks that form general higher-order complexes. We find that the same autoregulation rule applies across interaction complexity, provided that (1) interactions remain reversible and (2) complexes remain dilute rather than forming dense liquid phases. For liquid-liquid phase separation, the rule may still drive learning but is not mathematically guaranteed to be optimal; we leave this case to future work.

Figure 5a demonstrates this generality by training two 10-species networks, one with only dimerization and one including trimerization. Equilibrium constants (*K*_*ij*_ and *K*_*ijk*_) were independently sampled from a log-normal distribution, and both networks were trained on the same input-output examples (red circles). Despite distinct interaction types, both networks learned the same input-output response, with the same training rule producing different trained values of [*H*^tot^].

**FIG. 5.**
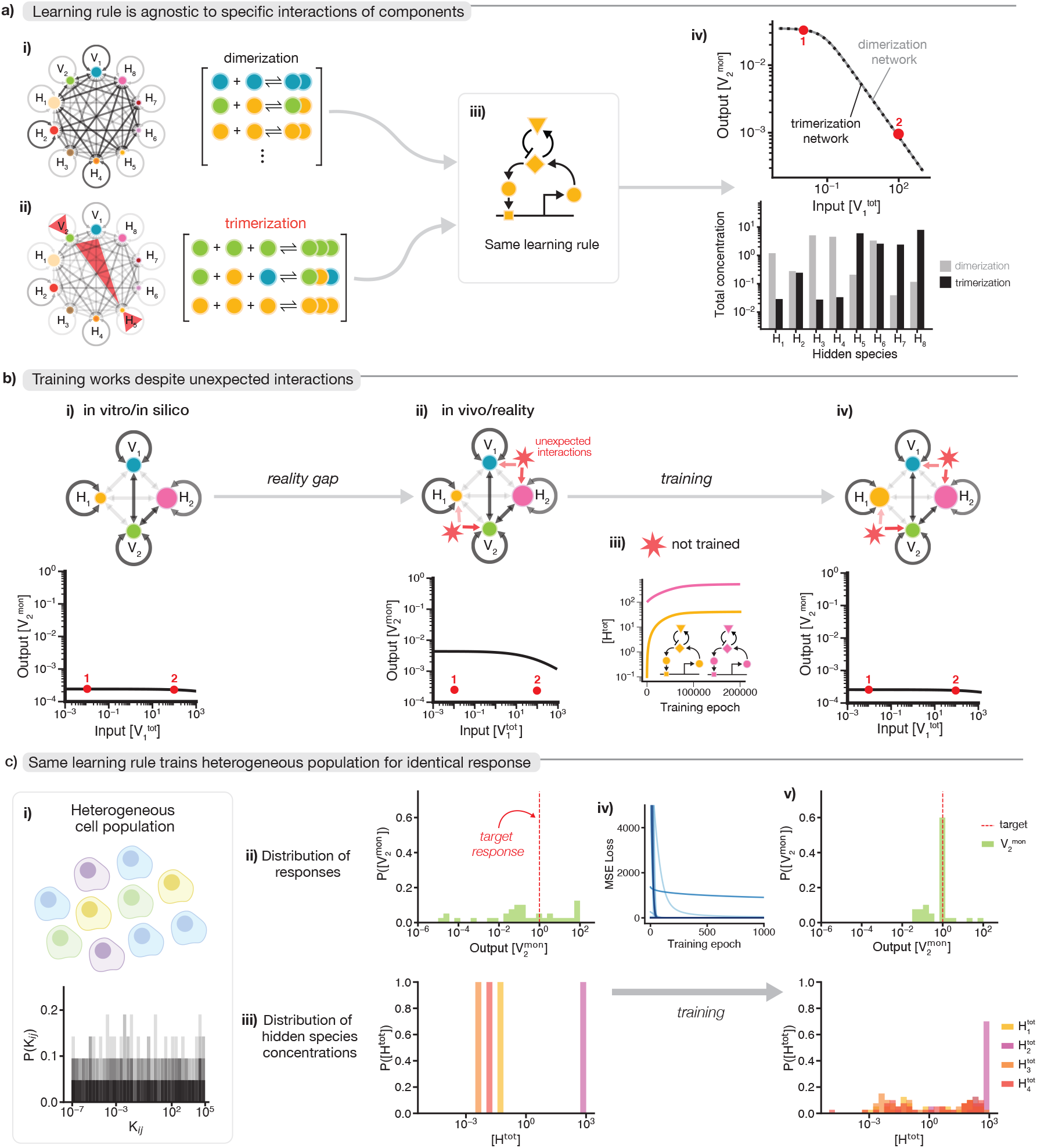
Auto-regulatory training is model-free and hence robust to unmodeled interactions and population heterogeneity. **a**. A ten-species network trained with either (i) trimerization or (ii) dimerization interactions. The same auto-regulatory training rule successfully trains both networks to identical input-output behavior using the same environmental examples, even though this requires tuning [*H*^tot^] to different values in the two networks. **b**. (i) A four-species network optimized *in silico* for a specific behavior produces (ii) incorrect behavior when unmodeled interactions with new interfering species (red stars) are introduced. (iii) Training via autoregulation on *H*_1_ and *H*_2_ alone recovers the desired input-output map without requiring knowledge, training or manipulation of the unmodeled species (red stars). **c**. (i) A heterogeneous population of cells, each with a 10-species network whose binding constants *K*_*ij*_ are randomly sampled from a wide distribution. (ii, iii) Initially, input-output behavior and hidden species concentrations vary widely across the population. (iv, v) After training in the same structured environment, each cell converges to different [*H*^tot^] values that compensate for that cell’s idiosyncratic *K*_*ij*_, so that all cells yield the same target response (red dashed line).

**FIG. 6.**
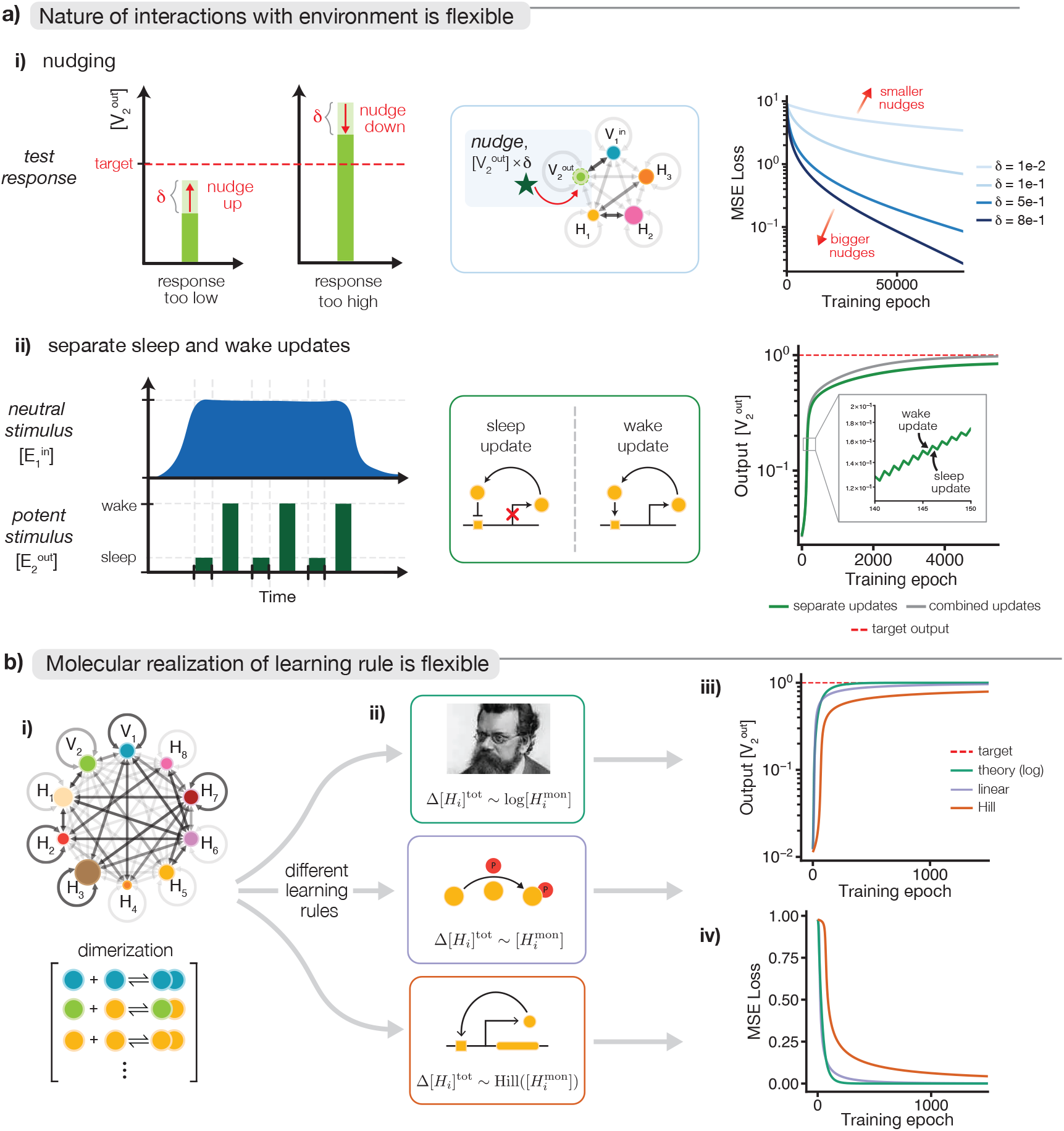
Robustness of the training framework to molecular details. **a**. Flexibility in training protocols: the training rule works with alternative models of environmental interaction. (i) The spiking environmental signal 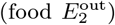 need only nudge the response molecule 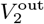 toward the desired level rather than imposing it exactly. A five-species network trained with nudging signals successfully decreases MSE loss, with larger nudges resulting in faster convergence. (ii) Separate sleep and wake phase auto-regulatory updates (with opposite signs) alleviate the need for rate-based autoregulation. Output 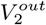 over sleep wake cycles converges to the target level (dashed red), but exhibits oscillations (inset) and larger final error relative to combined updates. **b**. (i, ii) Different mathematical forms of the auto-regulatory training rule (logarithmic, Hill, or linear proportional) applied to train a 10-species dimer network. (iii, iv) The mathematically optimal rule Δ[*H*^tot^] ∝ log[*H*^mon^], derived from Boltzmann machine theory, converges fastest, but molecularly realistic rules based on post-transcriptional activation (purple) or transcriptional autoregulation (orange) also successfully evolve toward the desired behavior.

This model-free property also addresses a central challenge in synthetic biology: the ‘reality gap’ between design and implementation. Synthetic circuits are typically designed from explicit or implicit models of molecular interactions, yet *in vivo* behavior can deviate from *in vitro* or *in silico* predictions due to unmodeled cellular interactions, forcing iterative redesign [77, 78].

Our feedback-based training framework bypasses this dependence on accurate models. In Fig. 5b, a four-species network is first designed *in silico* by choosing *K*_*ij*_ and [*H*^tot^] to match a target input-output response, but adding unmodeled interactions with additional cellular species (red stars) disrupts the behavior. Training the hidden species *H*_1_, *H*_2_ via autoregulation nevertheless restores the target response, without manipulating the interfering species (red stars) and without modifying the training rule to account for them.

Figure 5c extends this feature to heterogeneous populations. We simulated a population of 50 cells with 10-species networks in which each cell’s binding constants *K*_*ij*_ vary across orders of magnitude. Before training, a shared [*H*^tot^] produces diverse responses across cells; after exposure to the same training protocol, the auto-regulatory rule drives each cell to a different [*H*^tot^] that compensates for its specific *K*_*ij*_. Consequently, all cells achieve the same target output (red dashed line), showing that a single training environment can program diverse molecular systems to a common function.

## V. THE TRAINING FRAMEWORK IS ROBUST TO IMPLEMENTATION DETAILS

The model-free advantages above come with an apparent overhead: unlike static circuit design, training requires a plasticity mechanism (here, autoregulation of hidden-species levels) and an interaction protocol with the environment during a training period. A key practical question is how demanding those costs are in implementation. If successful training required a finely tuned, chemically specific realization of the training rule or a highly controlled training protocol, the added complexity could negate its benefits. Here we show that this is not the case: the framework’s performance is insensitive to many details of how training is realized, suggesting that the additional engineering burden can be modest.

First, training does not require the environment to clamp the output to its exact target. Figure 5e (top) shows that a simple “nudging” protocol, inspired by equilibrium propagation [79] and related physical implementations [80–82], is sufficient: after observing the output, the environment applies a small corrective push in the appropriate direction. Second, training can be discretized into separate sleep and wake phases with updates applied once per phase rather than continuously (Figure 5e, bottom). Although this slows learning, it relaxes the need for rate-based autoregulation and still converges to the desired behavior (see SI for more).

Finally, the precise functional form of autoregulation is flexible. While the Boltzmann-machine derivation (SI Sec. I) yields an optimal update Δ[*H*^tot^] ∝ log[*H*^mon^], Figure 5d shows that molecularly plausible alternatives also train successfully, including linear responses (used throughout this paper) that might model post-translational activation of hidden species and Hill-type nonlinearities typical of transcriptional feedback. These variants primarily affect convergence speed rather than whether learning succeeds.

Together, these results indicate that the framework’s benefits do not hinge on a narrow mechanistic implementation, and that the extra overhead of training can be realized with coarse, biophysically natural approximations rather than finely tuned control.

## VI. EXPERIMENTAL SIGNATURES

The most direct experimental test of the framework is constructive: build a molecular circuit with dense reversible interactions and rate-sensitive autoregulation of a subset of species, and ask whether it can be trained *in situ* even when embedded in a cellular background with unmodeled couplings.

In natural systems, the key prediction is cue substitution after paired experience of cues, and trained immunity provides a concrete setting for this test. Monocytes and macrophages can acquire durable changes in later responsiveness after transient exposure to stimuli such as *β*-glucan or BCG, with memory stored in metabolic and chromatin states [19]. Our framework suggests asking whether such memory can be extended from single-cue history to cue correlations. For example, a weak tissue- or tumor-B signal *X* could be repeatedly paired with *β*-glucan or BCG (the potent *Y*). After rest, the decisive test is whether *X* alone now evokes a trained-macrophage state as measured by cytokine output.

Cancer drug adaptation provides another setting. Drug-induced stress can establish drug-tolerant states through chromatin remodeling by promiscuous transcription-factor hubs such as AP-1 [83, 84]. If a neutral environmental cue (hypoxia, matrix stress) is repeatedly paired with a transient drug exposure, the cue alone should later preactivate a drug-adapted state and increase survival on rechallenge. More broadly, analogous paired-cue tests could be designed in fibroblast mechano-inflammatory memory [85, 86] or *Stentor* habituation, where slow memory-like variables and accessible readouts are already established [15, 20].

A specific molecular prediction of our framework is that longer-term learning should be visible in the short-term fluctuations of candidate training variables. Species *H*_*i*_ with the largest short-timescale fluctuations in their monomer-multimer distribution (as the environment enters and leaves the paired *X* + *Y* state) should show the largest long-timescale drift in 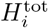, in the direction that dampens those fluctuations. This cross-timescale signature also addresses a recurring challenge in experimental studies of cellular learning, where many molecular species respond to environmental changes but most merely report the current stimulus rather than encode historical correlations. Species that fluctuate during training but show no directional drift in their total concentrations are stimulus-response variables rather than training variables.

Although our minimal model stores learned information in hidden-species total concentrations, natural systems may use such concentration changes as a proximal training layer that is later consolidated into longer-lived substrates such as chromatin. We therefore predict that persistent chromatin changes should accumulate preferentially at loci controlled by factors whose monomer-multimer distribution changes sharply during the *X* + *Y* training transition. After training, *X* alone should partially reactivate this chromatin-associated program, while perturbing the transient factors or chromatin writers during training should impair consolidation without necessarily blocking the immediate response to *Y*.

Finally, fast-onset, slow-offset (*τ*_*f*_ ≪ *τ*_*s*_) *Y* pulses should train more effectively than symmetric ramps (*τ*_*f*_ ≈ *τ*_*s*_) with the same *X*–*Y* correlation and total exposure, because the update rule depends on rapid transitions followed by slow relaxation. This control would distinguish the proposed mechanism from generic priming or learning rules that depend only on cue co-occurrence.

## VII. DISCUSSION

We showed that dense, reversible molecular interaction networks can be trained by rate-sensitive autoregulation of hidden-species concentrations, supporting Pavlovian conditioning, supervised classification, and tuning of stochastic phenotypic fractions within a lifetime. More broadly, this work demonstrates that physical systems can naturally embody learning algorithms typically associated with neural networks, in line with learning in electrical, mechanical, and metamaterial networks [79– 81, 87–101]. The key molecular ingredients, promiscuous interactions, autoregulation, and rate-sensitive signaling, are widespread in biology, suggesting that cells in structured environments could tune their responses to environmental statistics on timescales much shorter than evolution. Possible natural contexts include trained immunity, inflammatory memory, and non-genetic cancer drug resistance, where repeated environmental or therapeutic exposures can drive adaptive chromatin and regulatory remodeling; AP-1-dependent stress responses provide one candidate molecular substrate [83, 84].

Our results also suggest a model-free paradigm for synthetic biology. Traditional circuit design relies on detailed mechanistic models and repeated rounds of re-design when in vivo implementations deviate from expectations. By contrast, a trainable circuit can be assembled from expressive but imperfect components and trained directly *in situ* [56, 57], so that molecular weights self-adjust to realize a desired function despite unknown interactions. For example, an engineered immune cell could be trained on panels of healthy and cancer cells until it reliably triggers an effector program, exploiting real correlations in the training environment rather than a predefined model.

Our analysis has important limitations that suggest directions for future work. The auto-regulatory training rule was derived for reversible binding and assumes a separation of timescales between molecular equilibration, autoregulation, and environmental variation, and was limited to static input–output tasks. As a consequence, dense condensates [62] and complex temporal sequence processing [29] fall outside the present analysis. Another key challenge involves long term inheritance of learned behavior across many cell divisions. Persistent memory of learned hidden-species expression levels is likely to require multistable attractors for those species, epigenetic memory [102, 103], or continual reinforcement by the environment; we discuss these models in SI Sec.VI but need to be explored further in future work. Future work should alsop probe how evolution and learning processes might interact and sculpt trainable networks, for example by evolving network architectures under selection for rapid, flexible trainability across diverse tasks within a lifetime.

## VIII. ACKNOWLEDGEMENTS

We thank Lulu Qian, Erik Winfree, Cameron Chalk, Deborah Fygenson, Sidney Nagel, Pepijn Moerman, Max Schelling and members of the Murugan lab for discussions. KT acknowledges support from the U.S. National Science Foundation Graduate Research Fellowship under grant no. 2140001. MJF is supported in part by the Eric and Wendy Schmidt AI in Science Fellowship, a program of Schmidt Sciences. AM acknowledges support from the NIGMS of the National Institutes of Health under award number R35GM151211 and the National Science Foundation through DMR-2239801. M.B.E. is a Howard Hughes Medical Institute investigator, and was also supported by the Alfred P. Sloan Foundation (award number G-2024-22436) and the Chan Zuckerberg Initiative (award number 2024-349887). This work was supported by the University of Chicago’s Research Computing Center for computing resources, the National Science Foundation through the Center for Living Systems (PHY-2317138), and the NSF-Simons National Institute for Mathematics and Theory in Biology (National Science Foundation award DMS-2235451 and Simons Foundation award MPS-NITMB-00005320).

