## Supplemental Information for "Trainable computation in molecular networks"

### CONTENTS

Outline of the Supplementary Information 2

I. Sleep-wake learning theory in molecular systems 2
A. Sleep-wake training rules for general statistical systems 3
B. Specializing to molecular complexation networks 4

II. How the environment couples to visible species 5
A. Associative and targeted coupling 5
1. Associative training 5
2. Targeted training 6
B. Molecular mechanisms 6
1. Receptor-mediated model example 6
2. Coupling to monomeric concentrations 7
3. Coupling to total concentrations 7
C. A continuum of environmental coupling strengths 8
D. Verifying clampability 8

III. Minimal procedure for training 8
A. Monomer training for dimerization networks 9
B. Convergence in a two-species example 9

IV. Unsupervised learning in multi-stable networks 10
A. Toy 1D model 10
B. Positive autoregulation example 11
C. Gene toggle switch example 11

V. High-pass filtering mechanisms across biology 12
A. Temporal contrastive learning theory 12
B. Molecular realizations 13
1. Integral Feedback Control 13
2. Incoherent Feedforward Loops 14
3. Receptor Desensitization (Conformational Switching) 14
4. Phase Separation 15

VI. Timescales for learning and long-term memory 15
A. Timescales in molecular sleep-wake 15
B. Long-term memory of learned behavior 15

VII. Computational details 16
A. Simulating networks 16
1. Modeling dimerization 16
2. Parameters 17
3. Packages 17
B. Training details 17
1. Training protocol 17
2. Clamping concentrations 18
C. Choosing training targets 18
D. Figure descriptions 18
1. Pavlovian conditioning 18
2. Step-down to step-up 18
3. Monotonic to non-monotonic 18
4. Higher dimensional training 19
5. Iris classification 19
6. Generative learning 19
7. Trimerization networks 20
8. Networks with unmodeled interactions 20
9. Learning in heterogeneous populations 20

10. Alternate clamping rules 20
11. Alternate update dynamics 21
12. Alternate training rules 21

References 21

### OUTLINE OF THE SUPPLEMENTARY INFORMATION

This Supplementary Information provides the (a) theoretical derivations, (b) biological mechanisms, and (c) computational methods supporting the main text. The guide below explains where these materials are presented.

*a. How does this work biologically?* Readers interested primarily in the molecular mechanisms and their biological plausibility can focus on: Sec. II B (how environmental signals couple to visible species, including concrete molecular realizations such as set-point circuits and sequestration), Sec. V (biological mechanisms such as integral feedback, incoherent feedforward loops, receptor desensitization, phase separation that implement the required high-pass filtering), and Sec. VI (timescale requirements for learning and strategies for long-term memory, including epigenetic mechanisms in dividing cells).

*b. How is this justified mathematically?* Readers seeking the theoretical foundations can follow: Sec. I (derivation of the sleep-wake training rule for general statistical systems, specialization to molecular complexation networks), Sec. III (description and proof of convergence for a minimal training procedure), Sec. IV (extension of the framework to multi-stable networks for unsupervised/generative learning), and Sec. V (Sec. V A: how temporal contrastive learning resolves the temporal non-locality of sleep-wake updates without explicit memory storage).

*c. What was done in the simulations?* Sec. VII contains all computational details: network models and parameters, training protocols, clamping procedures, and figure-by-figure descriptions of every simulation in the main text.

For reference, the sections are organized as follows:

1. **Sleep-wake learning theory** (Sec. I): Derivation of the training rule from Boltzmann-machine principles and specialization to molecular complexation networks of arbitrary order.

2. **How the environment couples to visible species** (Sec. II B): Targeted vs. associative learning, molecular mechanisms of coupling, and verification of the clampability condition.

3. **A minimal procedure for training networks** (Sec. III): A minimal, concrete proposal to allow for training targeted monomer outputs in dimerization networks and a convergence proof for this example.

4. **Unsupervised learning in multi-stable networks** (Sec. IV): Extension to stochastic, multi-stable systems that can learn distributions over phenotypic states.

5. **High-pass filtering mechanisms** (Sec. V): Theory of temporal contrastive learning and a catalog of biological mechanisms that naturally implement high-pass filtering.

6. **Timescales and long-term memory** (Sec. VI): Timescale ordering required for learning, and strategies for maintaining learned behavior across cell divisions.

7. **Computational details** (Sec. VII): Full simulation parameters, training protocols, and figure-by-figure descriptions.

### I. SLEEP-WAKE LEARNING THEORY IN MOLECULAR SYSTEMS

The theoretical justification for our proposed single-cell learning framework is based in sleep-wake contrastive learning. A key feature of sleep-wake's appeal is its locality; updates to a unit $i$ only require information that was physically accessible to unit $i$.

In particular, sleep-wake algorithms were developed in the context of Boltzmann machines [1] – neuronal systems described as a set of visible neuron nodes $\mathcal{V} = \{V_1, ..., V_k\}$, coupled to a set of hidden nodes $\mathcal{H} = \{H_1, ..., H_k\}$ via an energy function $E_\lambda(\mathcal{V}, \mathcal{H})$ that the neurons collectively minimize. $E$ is in turn parameterized by a set of weights $\lambda$. For fixed weights $\lambda$, the values of the hidden nodes $\mathcal{H}$ sample distributions that are localized to the minima of $E_\lambda(\mathcal{V}, \mathcal{H})$. The goal of sleep-wake algorithms is to update $\lambda$ to $\lambda'$ such that new minima in $E_{\lambda'}(\mathcal{V}, \mathcal{H})$ correspond to samples of some target distribution. Spontaneous sampling of $E_{\lambda'}(\mathcal{V},\mathcal{H})$ can then serve as a generative model of the target distribution.

To achieve the correct update of weights $\lambda$, training alternates between two phases: sleep and wake. In the sleep phase, the system is decoupled from the environment and excitations of $\mathcal{V}$ and $\mathcal{H}$ are simply set by spontaneous sampling of $E_\lambda(\mathcal{V},\mathcal{H})$ from a Boltzmann distribution: $P(\mathcal{H},\mathcal{V}) \sim e^{-E_\lambda(\mathcal{H},\mathcal{V})}$. In the wake phase, the system experiences its environment, which sets a distribution over the values of the visible nodes $\mathcal{V} \sim Q(v)$.

We will make the notion of sleep and wake states more precise in the following sections, but the overarching goal is to nudge the system into having its natural sleep phase look like the wake phase. Therefore, weights $\lambda$ are updated to reinforce correlations in the wake phase in a Hebbian manner and penalize correlations in the sleep phase in an anti-Hebbian manner using the following training rule:

$$\frac{d\lambda_i}{dt} \propto \left\langle \frac{\partial E}{\partial \lambda_i} \right\rangle_{\text{sleep}} - \left\langle \frac{\partial E}{\partial \lambda_i} \right\rangle_{\text{wake}}. \tag{1}$$

This training procedure has several appealing features. First, the procedure naturally terminates once the sleep and wake phases become identical. Second, the procedure is usually local; for standard energy functions of physical systems, updates to $\lambda_i$ only involve information that involves the index $i$, or terms in the energy that couple to index $i$.

However, this locality in space is accompanied by a non-locality in time; the sleep and wake states have to be experienced at different times. For molecular or other physical systems, this temporal non-locality is an issue. One possibility to overcome temporal non-locality is to employ temporal contrastive learning (TCL) [2]. In this framework, weights update according to

$$\frac{d\lambda_i}{dt} \propto F_{HP}\left(\frac{\partial E}{\partial \lambda_i}\right) , \tag{2}$$

as the environment dynamically shifts between sleep and wake distributions. For temporal contrastive learning, $F_{HP}$ is any function which acts as a high-pass filter. We will return to the issue of how molecular systems can achieve sleep-wake updates without temporal storage in Sec. V.

Putting aside the issue of temporal non-locality for now, we will derive Eq. 1 for networks of molecules that form complexes through a dense network of interactions.

#### A. Sleep-wake training rules for general statistical systems

We derive a Boltzmann-machine training rule for general statistical systems and then specialize it to molecular networks. Related derivations were first given for chemical reaction networks in [3] and extended in [4]. Our emphasis is a formulation suited to the thermodynamic limit, where sleep and wake ensembles can have very different typical macrostates, yet remain connected by a physically minimal intervention we refer to as *clamping.* This leads to a *clampability* condition that must hold between the sleep and wake distributions if learning is to proceed with minimal additional intervention.

Let a physical system sample microstates $\sigma$ from $P_\lambda(\sigma) = e^{-G_\lambda(\sigma)}/Z_\lambda$, with $Z_\lambda = \sum_\sigma e^{-G_\lambda(\sigma)}$, where $\lambda$ denotes arbitrary trainable parameters. Let $v = \nu(\sigma)$ be an observable, inducing $P_\lambda(v) = \sum_{\sigma:\,\nu(\sigma)=v} P_\lambda(\sigma)$. Given a target distribution $Q(v)$, we choose $\lambda$ so that $P_\lambda(v)$ approximates $Q(v)$ by minimizing

$$D_{KL}(Q||P_\lambda) = \sum_v Q(v) \log \frac{Q(v)}{P_\lambda(v)}. \tag{3}$$

Differentiating with respect to $\lambda$ and using $\partial_\lambda \log P_\lambda(\sigma) = -\partial_\lambda G_\lambda(\sigma) + \langle \partial_\lambda G_\lambda \rangle_{P_\lambda}$ gives

$$\partial_\lambda D_{KL}(Q||P_\lambda) = -\left\langle \partial_\lambda G_\lambda(\sigma) \right\rangle_{P_\lambda(\sigma)} + \sum_v Q(v) \left\langle \partial_\lambda G_\lambda(\sigma) \right\rangle_{P_\lambda(\sigma|v)}, \tag{4}$$

where $P_\lambda(\sigma|v) = P_\lambda(\sigma)\,\mathbf{1}\{\nu(\sigma) = v\}/P_\lambda(v)$ is the conditional. The first term is an expectation under the system's natural sampling (the *sleep* phase). The second averages the same quantity over microstates conditioned on a target distribution over $v$.

**Clampability.** We assume there is a physical procedure that enforces the desired distribution over $v$ while leaving all remaining degrees of freedom distributed as the system's conditional distribution. Concretely, we assume that the resulting *wake* distribution over microstates satisfies

$$Q(\sigma) = Q(v)\, P_\lambda(\sigma|v), \qquad v = \nu(\sigma). \tag{5}$$

This 'clampability' condition is a requirement for implementing learning with minimal additional engineering. Equivalently, $Q(\sigma)/P_\lambda(\sigma)$ depends only on $v$ (up to normalization), so microstates $\sigma_1, \sigma_2$ with the same $v$ retain the same relative probability in sleep and wake. Note that while $Q(v)$ is fixed, $Q(\sigma)$ generally inherits $\lambda$-dependence through the conditional $P_\lambda(\sigma|v)$.

Under clampability (5), the second term in Eq. (4) becomes $\langle \partial_\lambda G_\lambda(\sigma)\rangle_{Q(\sigma)}$, yielding

$$\partial_\lambda D_{KL}(Q||P_\lambda) = \langle \partial_\lambda G_\lambda(\sigma)\rangle_{\text{wake}} - \langle \partial_\lambda G_\lambda(\sigma)\rangle_{\text{sleep}}\,, \tag{6}$$

and gradient descent recovers the sleep-wake training rule first expressed in Eq. 1:

$$\frac{d\lambda}{dt} = -\eta\, \partial_\lambda D_{KL}(Q||P_\lambda) = -\eta \left( \langle \partial_\lambda G_\lambda(\sigma)\rangle_{\text{wake}} - \langle \partial_\lambda G_\lambda(\sigma)\rangle_{\text{sleep}} \right). \tag{7}$$

#### B. Specializing to molecular complexation networks

We now specialize the general framework to molecular complexation networks. Consider a set of monomer species $\{m_i\}_{i=1}^N$ that can form dimers $\{d_{ij}\}_{i\le j}$ through reactions $m_i + m_j \to d_{ij}$, subject to mass conservation $m_i^{\text{tot}} = m_i + 2d_{ii} + \sum_{j\neq i} d_{ij}$.

Following Sec. I A, we work with discrete microstates $\sigma = \{m_1, \ldots, m_N, d_{11}, \ldots, d_{NN}\}$ specified by integer counts of each species. The microscopic free energy is

$$\mathcal{G}(\sigma) = \sum_i \mu_i\, m_i + \sum_{i\le j} \mu_{ij}\, d_{ij} + \sum_i \lambda_i^{\text{tot}} \big(m_i^{*,\text{tot}} - m_i^{\text{tot}}\big)\,, \tag{8}$$

where the Lagrange multipliers $\lambda_i^{\text{tot}}$ enforce mass conservation at the equilibrium values $m_i^{*,\text{tot}}$ that minimize $\mathcal{G}$, and the system samples from $P(\sigma) = e^{-\mathcal{G}(\sigma)}/Z$. Our goal is to adjust the conserved totals $m_i^{*,\text{tot}}$ to minimize $D_{KL}(Q\|P)$ via Eq. 6.

The monomer chemical potentials decompose as $\mu_i = \mu_i^{\text{ext}} + E_i + \ln(m_i/\tilde{V})$, where $E_i$ is the intrinsic formation energy, $\mu_i^{\text{ext}}$ is an externally imposed potential, and $\tilde{V}$ is the system volume. Dimer potentials take the analogous form $\mu_{ij} = \mu_{ij}^{\text{ext}} + E_{ij} + \ln(d_{ij}/\tilde{V})$, with $E_{ij} = E_i + E_j + \Delta E_{ij}$ and $\Delta E_{ij}$ the binding free energy. Since $\mathcal{G}$ depends on $m_i^{*,\text{tot}}$ only through the constraint term, one immediately has $\partial\mathcal{G}/\partial m_i^{*,\text{tot}} = \lambda_i^{\text{tot}}$. To determine $\lambda_i^{\text{tot}}$, we impose stationarity of $\mathcal{G}$ at equilibrium. Setting $\partial\mathcal{G}/\partial m_i = 0$ and $\partial\mathcal{G}/\partial d_{ij} = 0$ yields

$$\lambda_i^{\text{tot}} = \mu_i^{\text{ext}} + E_i + \ln m_i^* + 1\;, \qquad \lambda_i^{\text{tot}} + \lambda_j^{\text{tot}} = \mu_{ij}^{\text{ext}} + E_{ij} + \ln d_{ij}^* + 1\;, \tag{9}$$

where $*$ denotes equilibrium values. Substituting the first relation into (6), the terms $E_i$, $\mu_i^{\text{ext}}$, and the additive constant are all properties of the molecular species and unchanged between sleep and wake, so they cancel. Only $\ln m_i^*$ varies between phases, yielding a training rule

$$\frac{d}{dt} m_i^{*,\text{tot}} \propto \langle \ln m_i^* \rangle_{\text{sleep}} - \langle \ln m_i^* \rangle_{\text{wake}}\;. \tag{10}$$

**Thermodynamic limit.** Equivalently, one can work directly in terms of number concentrations (denoted by brackets). In the dilute-solution limit, the Gibbs free energy at fixed temperature and pressure (in units of $k_BT$, normalized by the number of solvent molecules $n_s$) takes the form

$$\mathcal{G} = \sum_i [m_i^{\text{mon}}](\mu_i^\circ + \ln[m_i^{\text{mon}}] - 1) + \sum_{i\le j} [d_{ij}](\mu_{ij}^\circ + \ln[d_{ij}] - 1) + \sum_i \lambda_i^{\text{tot}}\big([m_i^{*,\text{tot}}] - [m_i^{\text{tot}}]\big)\;, \tag{11}$$

where $\mu_\alpha^\circ$ denotes the standard chemical potential of species $\alpha$, and the last term stems from the constraint that we satisfy mass conservation $[m_i^{*,\text{tot}}] = [m_i^{\text{tot}}] \equiv [m_i^{\text{mon}}] + 2[d_{ii}] + \sum_{j\neq i}[d_{ij}]$. As before, one finds $\partial\mathcal{G}/\partial[m_i^{*,\text{tot}}] = \lambda_i^{\text{tot}}$, which from stationarity of $\mathcal{G}$ with respect to $[m_i^{\rm mon}]$ implies $\partial\mathcal{G}/\partial[m_i^{*,\rm tot}] = \ln[m_i^{\rm mon}] + \mu_i^\circ$. Applying Eq. 6, the training rule for total monomer concentrations is therefore

$$\frac{d}{dt}[m_i^{*,\rm tot}] \propto \langle \ln{[m_i^{\rm mon}]} \rangle_{\rm sleep} - \langle \ln{[m_i^{\rm mon}]} \rangle_{\rm wake} \ , \tag{12}$$

i.e., total amounts are adjusted based on the difference in log-monomer concentrations between sleep and wake phases. As a corollary, since homodimer concentrations satisfy $[d_{ii}] = K_{ii}[m_i^{\rm mon}]^2$, where $K_{ii}$ is the equilibrium constant of homodimerization for species $m_i$, an equivalent training rule can be formulated as

$$\frac{d}{dt}[m_i^{*,\rm tot}] \propto \langle \ln{[d_{ii}]} \rangle_{\rm sleep} - \langle \ln{[d_{ii}]} \rangle_{\rm wake} \ . \tag{13}$$

**Extension to higher-order complexes.** This derivation extends straightforwardly to networks with higher-order complexes (trimers, tetramers, etc.). Writing the free energy as $\mathcal{G} = \sum_\alpha [n_\alpha](\mu_\alpha^\circ + \ln[n_\alpha] - 1)$ with mass conservation $[m_i^{*,\rm tot}] = [m_i^{\rm mon}] + \sum_\alpha s_\alpha^i [n_\alpha]$ (where $s_\alpha^i$ counts the copies of monomer $i$ in complex $\alpha$), the same calculation gives $\partial\mathcal{G}/\partial[m_i^{*,\rm tot}] = \ln[m_i^{\rm mon}] + \mu_i^\circ$. The monomer-based training rule Eq. (12) therefore holds unchanged.

**Robustness to unmodeled interactions.** The training rule also remains valid in the presence of unmodeled interactions. Additional dimerization partners or higher-order complexes involving unknown species enter mass conservation but do not alter the form of $\partial\mathcal{G}/\partial[m_i^{*,\rm tot}]$, since $[m_i^{\rm mon}] = [m_i^{\rm tot}] - (\text{all complexes containing } i)$ regardless of the identity of those complexes. Hence learning updates to $[m_i^{*,\rm tot}]$ remain valid in the dilute limit even with unknown interactions present.

### II. HOW THE ENVIRONMENT COUPLES TO VISIBLE SPECIES

#### A. Associative and targeted coupling

Even with the same internal training dynamics described in Sec. I, the mode of learning depends on how the environment couples to visible species. In this section, we will discuss two modes – associative and targeted – which make different demands on the type of network-environment coupling and lead the networks to solve different learning problems.

##### 1. *Associative training*

In associative training, the environment presents correlated molecular signals that couple strongly enough to visible species to override their internal equilibrium. Crucially, the environment does not prescribe what the internal response of the network — the values that visible species take on during the wake phase — should be; the response is simply whatever the environment naturally induces.

As a concrete example, consider a cell with a "bell" input species coupled to environmental signal $E_B$ and a "food" response species whose monomer concentration is $V_F^{\rm mon}$. In associative training, the environment presents correlated pairs $(E_B, E_F)$ drawn from a distribution $P(E_B, E_F)$, and the resulting internal response $V_F^{\rm mon}$ is simply whatever the network produces given the environmental signals.

Formally, if we denote the steady-state response as $V_F^{\rm mon}(E_B,\, E_F)$, which is independent of $\{H_i^{\rm tot}\}$, then the goal of associative training is

$$V_F^{\rm mon}\big(E_B, 0; \{H_i^{\rm tot}\}\big) \ \approx \ V_F^{\rm mon}\big(E_B, E_F^*\big), \tag{14}$$

where $E_F^*$ is the food level that co-occurs with $E_B$ in the environment. The "target" response on the right-hand side is determined only by $E_F^*$ — the cell is not learning toward a fixed internal response, but toward reproducing its own environmentally-shaped response. When the coupling between the environment and the cell interior is strong, the steady-state response depends only on the environmental signals and not on the current hidden-species state. Training succeeds when presenting $E_B$ alone produces the same internal state as would have been produced by the joint presentation of $(E_B, E_F)$.

The training procedure is straightforward: the environment simply provides correlated signals, and the autoregulatory learning rule does the rest. No feedback control or monitoring of the response variable is required. This mode is natural when one views the cell as passively experiencing a structured environment, internalizing environmental correlations such that partial external cues reconstruct the full internal response. This is the mode used for Pavlovian conditioning in the main text (Sec. II).

#### 2. Targeted training

In targeted training, the environment (or an experimenter) demands that a *specific* internal response $r^*$ be produced. The goal is

$$V_F^{\rm mon}\big(E_B, 0; \{H_i^{\rm tot}\}\big) \;\approx\; r^*, \tag{15}$$

where $r^*$ is a desired target response specified externally, not by the environmental food signal.

To achieve this using the same autoregulatory training rule, during each training cycle the environmental signal $E_F$ must be set to whatever value is needed to drive the response variable $V_F^{\rm mon}$ to $r^*$:

$$V_F^{\rm mon}\big(E_B,\, E_F;\, \{H_i^{\rm tot}\}\big) \;=\; r^*. \tag{16}$$

This inversion — setting $E_F$ to achieve a desired $V_F^{\rm mon}$ — requires some form of feedback control during training. The required $E_F$ may change over the course of training as $\{H_i^{\rm tot}\}$ evolve. Unlike associative training, the target $r^*$ is fixed externally.

Targeted training is more demanding (it requires feedback mechanisms), but it enables the cell to learn arbitrary input-output mappings specified by the environment. This mode is the relevant one for supervised learning of specific input-output mappings (main text Sec. III). Targeted training is also used for unsupervised (generative) learning of distributions (main text Sec. IV): during the wake phase, visible-species concentrations are fixed to samples drawn from a target distribution $Q([V_1^{\rm tot}], \ldots, [V_n^{\rm tot}])$, and the autoregulatory learning rule adjusts hidden-species levels to match the system's spontaneous distribution to $Q$.

An alternative way to achieve targeted training is *nudging*: rather than precisely enforcing $V_F^{\rm mon} = r^*$, one merely pushes the response variable in the direction of $r^*$, increasing $E_F$ slightly if the current response is below target and decreasing it if above. Nudging does not require exact feedback control but still achieves targeted training over many cycles, as shown by work on Equilibrium Propagation [5]. This is explored in the main text (Fig. 5).

### B. Molecular mechanisms

We now describe molecular mechanisms to realize the cell-environment coupling. Each mechanism may be more or less useful for associative or targeted training, depending on physical parameters.

#### 1. Receptor-mediated model example

We begin by introducing a concrete receptor-mediated model of strong environmental coupling to monomer concentrations, which we also present in Sec. II of the main text. We discuss how it may be used for both associative and targeted training.

In this model, an environmental ligand $E_k$ activates a pool of receptors, $R$ to $R^*$. The active receptor then catalyzes the transformation of $U$, available in effectively unlimited amounts, into visible species $V_k$. $V_k$ reversibly dimerizes with hidden species, $H_i$. To summarize,

$$R \xrightarrow{E_k} R^* \;, \qquad U + R^* \rightleftharpoons V_k + R^* \;, \qquad V_k + H_i \rightleftharpoons VH_{ki} \;.$$

In this model, the dynamics of $V_k^{\rm mon}$ are described by

$$\frac{d[V_k^{\rm mon}]}{dt} = f([E_k])[U] - \lambda([E_k])[V_k^{\rm mon}] + k_{\rm off}[VH] - k_{\rm on}[V_k^{\rm mon}][H^{\rm mon}] \;,$$

$$\frac{d[H^{\rm mon}]}{dt} = k_{\rm off}[VH] - k_{\rm on}[V_k^{\rm mon}][H^{\rm mon}] \;,$$

with conservation law $[H^{\rm tot}] = [H^{\rm mon}] + [VH]$.

*a. Wake phase:* Setting $\frac{d[V_k^{\rm mon}]}{dt} = 0$ with $E_k \neq 0$, and taking the strong-coupling limit $f([E_k])[U], \lambda([E_k]) \gg k_{\rm on}, k_{\rm off}$, the reversible binding terms become negligible and the fixed point is

$$[V_k^{\rm mon}]^* \approx \frac{f([E_k])[U]}{\lambda([E_k])} \;,$$

which, importantly, is independent of the training parameter $[H^{\rm tot}]$.

*b. Sleep phase:* When $[E_k] \to 0$, production and degradation of $V_k$ both vanish, and $[V_k^{\rm tot}]$ is conserved at its wake phase value. The fixed point is now determined solely by the binding equilibrium and the conservation laws for both $[H^{\rm tot}]$ and $[V_k^{\rm tot}]$, making $[V_k^{\rm mon}]^*$ a function of $k_{\rm off}/k_{\rm on}$ and $[H^{\rm tot}]$. In particular, increasing $[H^{\rm tot}]$ sequesters more $V_k$ into dimers, reducing $[V_k^{\rm mon}]^*$.

Effectively, in the wake phase, large $[E_k]$ pins $[V_k^{\rm mon}]^*$ to a value set by the environmental dynamics, independent of the binding equilibrium with $H_i$. Reducing $[E_k]$ toward zero in the sleep phase removes this environmental drive, leaving $[V_k^{\rm mon}]^*$ determined solely by $[H_i^{\rm tot}]$. In this way, $[E_k]$ naturally decides whether the environment or the internal binding dynamics control $[V_k^{\rm mon}]^*$. Training dynamics on $H_i$ can thus tune $[V_k^{\rm mon}]^*_{\rm sleep}$ to match the wake phase target.

Training in the associative mode would reconfigure the network to a new equilibrium where it assumes the internal response $[V_k^{\rm mon}]^*$ which was naturally induced by $E_k$, even in its absence. This model could also be used for targeted training, where one explicitly demands a specific response $[V_k^{\rm mon}]^* = r^*$. To accomplish this task, one would need to identify the particular value of $E_k^*$ which gives the target response $[V_k^{\rm mon}]^* \approx \frac{f([E_k^*])[U]}{\lambda([E_k^*])} = r^*$ and train the network with that concentration.

#### 2. *Coupling to monomeric concentrations*

In this class of mechanisms, an environmental molecule $E_k$ influences the monomer concentration $[V_k^{\rm mon}]$ of a visible species in a dimerization network (Sec. I B) with equilibrium constants $K_{ij}$. Because monomers are shared among dimers,

$$[V_k^{\rm tot}] = [V_k^{\rm mon}] + 2[V_{kk}] + \sum_{j\neq k}[V_{jk}] \ , \qquad [V_{jk}] = K_{jk}[V_k^{\rm mon}][V_j^{\rm mon}] \ ,$$

any mechanism that influences $[V_k^{\rm mon}]$ also implicitly affects $[V_k^{\rm tot}]$, which adjusts to satisfy equilibrium.

*a. Set-point circuit.* A biologically plausible mechanism is a *set-point circuit* in which $E_k$ modulates a production–degradation balance for $V_k^{\rm mon}$, e.g. through gene regulatory dynamics:

$$\frac{d}{dt}[V_k^{\rm mon}] = \kappa\frac{[V_k^{\rm mon}]}{[V_k^{\rm mon}] + K_D} - \lambda[V_k^{\rm mon}], \tag{17}$$

where $E_k$ controls one or more of $\kappa$, $K_D$, or $\lambda$, setting the steady-state monomer to $[V_k^{\rm mon}] = f([E_k])$. At infinite feedback gain, $[V_k^{\rm mon}]$ is pinned exactly (targeted). At finite gain, the circuit cannot fully compensate for the drain of monomers into dimers with hidden species, and $[V_k^{\rm mon}]$ depends on $\{H_i^{\rm tot}\}$ (associative).

*b. Reservoir.* A statistical-physics abstraction of this mechanism is a *reservoir* of $V_k$ monomers whose chemical potential is set by $E_k$. An infinite reservoir pins $[V_k^{\rm mon}] = f([E_k])$ regardless of hidden-species levels. A finite reservoir can be partially depleted as hidden species pull $V_k$ into dimers, so $[V_k^{\rm mon}]$ acquires a dependence on $\{H_i^{\rm tot}\}$.

Note that homodimer concentrations $[V_{kk}] = K_{kk}[V_k^{\rm mon}]^2$ could serve as the coupled observable instead, since clamping $[V_{kk}]$ is equivalent to clamping $[V_k^{\rm mon}]$ up to a known monotonic transformation.

#### 3. *Coupling to total concentrations*

In this class of mechanisms, the environmental molecule $E_k$ influences the effective total concentration of $V_k$ available for dimerization,

$$[\tilde{V}_k^{\rm tot}] = [V_k^{\rm mon}] + 2[V_{kk}] + \sum_{j\neq k}[V_{jk}] \ .$$

Unlike the monomer-coupling mechanisms of the previous subsection, even in the strong-coupling limit, only the total is pinned by $E_k$. The monomer concentration $[V_k^{\rm mon}]$ is then determined by the equilibrium partitioning of $[\tilde{V}_k^{\rm tot}]$ among monomers and dimers, which depends on hidden-species levels through $[V_{jk}] = K_{jk}[V_k^{\rm mon}][V_j^{\rm mon}]$. Away from the strong-coupling limit, both total and monomer depend on hidden species.

*a. Activation.* Environmental molecules $E_k$ convert an inactive form $V_k^{(in)}$ into an active form $V_k^{(a)}$ that can participate in dimerization:

$$E_k + V_k^{(in)} \to V_k^{(a)}.$$

If driven to completion, $[\tilde{V}_k^{\text{tot}}] = [E_k]$ (targeted). Biochemical realizations include spatial localization (e.g. hormones providing a nuclear localization signal to cytoplasmic steroid receptors) and phosphorylation by receptor kinases. If the activation is incomplete or reversible, $[\tilde{V}_k^{\text{tot}}]$ depends on the network state.

*b. Sequestration.* Environmental molecules $E_k$ bind to and sequester visible monomers, removing them from the dimerization pool. The sequestered complex $EV$ is formed by mass action, $E_k + V_k^{\text{mon}} \rightleftharpoons EV$ with $[EV] = K_{EV}[E_k][V_k^{\text{mon}}]$, giving an effective total

$$[\tilde{V}_k^{\text{tot}}] = [V_k^{\text{tot}}] - [EV] = [V_k^{\text{mon}}] + 2[V_{kk}] + \sum_{j\neq k}[V_{jk}] \ .$$

In the limit of irreversible binding ($K_{EV} \to \infty$), $E_k$ removes a fixed number of molecules regardless of the network state, setting $[\tilde{V}_k^{\text{tot}}]$ to a value determined entirely by $[E_k]$ (targeted; Sec. II C). At finite $K_{EV}$, the amount sequestered depends on $[V_k^{\text{mon}}]$, which itself depends on hidden-species levels.

#### C. A continuum of environmental coupling strengths

The molecular realizations described in Sec. II B may be used for either associative or targeted training and can achieve a continuum of environmental-coupling strengths depending on physical parameters like reservoir size, feedback gain, and binding affinity. Associative training requires strong coupling, where the environment completely overwhelms the system's dynamics to determine the internal response. Targeted training, in contrast, is less demanding in this regard. The closed-feedback loop that monitors and adjusts $E_F$ to enforce a desired response $r^*$ compensates for the network's internal dynamics, enabling training at any coupling strength.

In this paper, we primarily work in the targeted limit, where the environment is assumed to set the visible-species concentrations to exact values independent of the hidden-species state. This simplifies the analysis and permits the supervised learning of specific input-output mappings and unsupervised learning explored in the main text.

#### D. Verifying clampability

As a final theoretical note, for the sleep-wake training rule to be valid, each coupling mechanism must satisfy the clampability condition in Eq. (5): for two microstates $\sigma_1, \sigma_2$ with the same value of the clamped observable, the environmental perturbation must leave their relative probability unchanged. We verify this for both monomer clamping and total-amount clamping.

In both cases, the argument has the same structure. The environmental coupling shifts a conjugate field in the free energy (8), adding a term of the form $\delta \cdot \nu(\sigma)$ to $\mathcal{G}(\sigma)$, where $\delta$ is the magnitude of the shift and $\nu(\sigma)$ is the clamped observable. Under this perturbation, the probability of a microstate becomes $P^*(\sigma) \propto e^{-\mathcal{G}(\sigma) - \delta\,\nu(\sigma)}$. For two microstates $\sigma_1, \sigma_2$ with the same $\nu(\sigma_1) = \nu(\sigma_2) = v$, the ratio

$$\frac{P^*(\sigma_1)}{P^*(\sigma_2)} = \frac{e^{-\mathcal{G}(\sigma_1) - \delta\, v}}{e^{-\mathcal{G}(\sigma_2) - \delta\, v}} = \frac{e^{-\mathcal{G}(\sigma_1)}}{e^{-\mathcal{G}(\sigma_2)}} = \frac{P(\sigma_1)}{P(\sigma_2)} \tag{18}$$

is unchanged, so clampability (5) holds. We now verify this for the coupling mechanisms introduced below.

**Monomer clamping.** Fixing $[V_k^{\text{mon}}]$ shifts the monomer chemical potential, $\mu_k \to \mu_k + \delta\mu_k$, adding $\delta\mu_k \cdot m_k$ to $\mathcal{G}(\sigma)$. This cancels at fixed $m_k$. Because only the monomer chemical potential shifts while dimer chemical potentials are unchanged, this perturbation effectively modifies the equilibrium constants $K_{kj}$.

**Total amount clamping.** Fixing $[V_k^{\text{tot}}]$ shifts the conservation-law multiplier, $\lambda_k^{\text{tot}} \to \lambda_k^{\text{tot}} + \delta\lambda_k^{\text{tot}}$, adding a term proportional to $m_k^{\text{tot}}$ to $\mathcal{G}(\sigma)$. This cancels at fixed $m_k^{\text{tot}}$. Here, the chemical potentials of all species containing $V_k$ shift by the same amount, so the equilibrium constants $K_{ij}$ are unchanged and only the conserved total $[V_k^{*,\text{tot}}]$ is modified.

### III. MINIMAL PROCEDURE FOR TRAINING

We now describe and justify a concrete training procedure for monomer outputs in a dimerization network and prove convergence in a two-species example.

#### A. Monomer training for dimerization networks

To achieve targeted monomer outputs in a dimerization network, we adopt the following minimal procedure: we first define a *sleep phase*, where the system equilibrates without environmental coupling, and all species concentrations are determined by mass conservation and equilibrium constants:

$$\begin{cases} m_i^{*,\mathrm{tot}} & = m_i + 2d_{ii} + \sum_{j\neq i} d_{ij} \\ K_{ij} & = \frac{d_{ij}}{m_i m_j} \end{cases} \tag{19}$$

In the *wake phase*, the environment fixes the monomer concentration of a visible output species $m_0$ to a target value $m_0^*$, while total amount of species $m_0$ is no longer constrained. However, hidden species $i > 0$ retain their mass conservation, and all equilibrium constants remain unchanged:

$$\begin{cases} m_i^{*,\mathrm{tot}} & = m_i + 2d_{ii} + \sum_{j\neq i} d_{ij} \quad \forall i > 0 \\ m_0 & = m_0^* \\ K_{ij} & = \frac{d_{ij}}{m_i m_j} \end{cases} \tag{20}$$

Hidden species $m_i$ with $i > 0$ then have total amounts updated as:

$$\frac{d}{dt} m_i^{*,\mathrm{tot}} \propto \langle \ln m_i^* \rangle_{\mathrm{sleep}} - \langle \ln m_i^* \rangle_{\mathrm{wake}} \ . \tag{21}$$

while $m_0^{*,\mathrm{tot}}$ remains fixed.

**Relation to the general training rule.** The sign of the update rule (21) is opposite to the general rule (12). The origin of the sign flip can be seen by noting that the sleep phase system of equations (19) clamps $m_0^{*,\mathrm{tot}}$ to be a set target value; meanwhile, $m_0^{*,\mathrm{tot}}$ is unconstrained in the wake phase equations (20). Hence our procedure can equivalently be viewed as total amount clamping (Sec. II D) with sleep and wake labels exchanged, and therefore training drives $m_0^{*,\mathrm{tot}}$ in the unconstrained phase toward its value in the constrained phase. As $m_0^{*,\mathrm{tot}}$ converges in value between the two phases, monomer concentration $m_0$ in the sleep phase also converges to the target $m_0^*$ enforced in the wake phase. To account for this redefinition of sleep and wake phases relative to the straightforward procedure of total amount clamping, we flip the sign of the update rule (21).

We now work through a simple two-species network to identify the training fixed point and show the update rule results in convergence to the fixed point.

#### B. Convergence in a two-species example

Consider monomers V and H forming a heterodimer VH with equilibrium constant $K_{VH}$. The goal is to train $H^{\mathrm{tot}}$ so that the equilibrium monomer concentration $V^{\mathrm{mon}}$ matches a target value $e^{\mu_V}$, with $V^{\mathrm{tot}}$ fixed.

**Wake phase.** In the wake phase, a fixed chemical potential $\mu_V$ externally sets the concentration of $V^{\mathrm{mon}}$ to $e^{\mu_V}$, while $H^{\mathrm{tot}}$ is conserved. The equilibrium concentrations are determined by

$$\begin{cases} H^{\mathrm{tot}} = H + VH \\ K_{VH} = \frac{VH}{(V)(H)} \\ V = e^{\mu_V} \end{cases}$$

which gives $H^{\mathrm{eq}}_{\mathrm{wake}} = H^{\mathrm{tot}}/(1 + e^{\mu_V} K_{VH})$.

**Sleep phase.** In the sleep phase, we replace the constraint on $V^{\mathrm{mon}}$ with mass conservation for $V^{\mathrm{tot}}$:

$$\begin{cases} H^{\mathrm{tot}} = H + VH \\ K_{VH} = \frac{VH}{(V)(H)} \\ V^{\mathrm{tot}} = V + VH \end{cases}$$

Eliminating $VH$ and $H$ yields a quadratic in $V^{\rm eq}_{\rm sleep}$, whose positive root gives a closed-form (but lengthy) expression for $H^{\rm eq}_{\rm sleep}$ as a function of $H^{\rm tot}$, $V^{\rm tot}$, and $K_{VH}$:

$$H^{\rm eq}_{\rm sleep} = \frac{1}{2}\left(H^{\rm tot} - \frac{1}{K_{VH}} - V^{\rm tot} + \sqrt{\frac{4K_{VH}V^{\rm tot} + [1 + K_{VH}(H^{\rm tot} - V^{\rm tot})]^2}{K_{VH}}}\right)$$

**Fixed point.** Training should terminate when there is no difference between sleep and wake, i.e., when $H^{\rm eq}_{\rm wake} = H^{\rm eq}_{\rm sleep}$. Equating the wake and sleep expressions for $H$ and solving for $H^{\rm tot}$ gives a unique fixed point at

$$H^{\rm tot*} = \frac{(1 + K_{VH}e^{\mu_V})(V^{\rm tot} - e^{\mu_V})}{K_{VH}e^{\mu_V}}, \qquad V^{\rm tot} > e^{\mu_V}. \tag{22}$$

Substituting $H^{\rm tot*}$ back into the sleep-phase equations confirms that $V^{\rm eq}_{\rm sleep} = e^{\mu_V}$, so the fixed point indeed achieves the desired target.

**Training rule and stability.** To drive the system toward this fixed point, the hidden-species total is updated according to

$$\frac{dH^{\rm tot}}{dt} \propto \langle \ln H \rangle_{\rm wake} - \langle \ln H \rangle_{\rm sleep}\ , \tag{23}$$

while $V^{\rm tot}$ remains fixed. Writing $\frac{dH^{\rm tot}}{dt} = f(H^{\rm tot})$, the fixed point is stable if $f'(H^{\rm tot*}) < 0$. Differentiating $f$ with respect to $H^{\rm tot}$ gives

$$\frac{d}{dH^{\rm tot}}\frac{dH^{\rm tot}}{dt} = \frac{1}{H^{\rm tot}} - \frac{K_{VH}[\Delta + 1 + K_{VH}(H^{\rm tot} - V^{\rm tot})]}{\Delta[\Delta - 1 + K_{VH}(H^{\rm tot} - V^{\rm tot})]}, \tag{24}$$

where $\Delta = \sqrt{([1 + K_{VH}(H^{\rm tot} - V^{\rm tot})]^2 + 4K_{VH}V^{\rm tot}}$.

Evaluating at $H^{\rm tot*}$using the closed-form sleep-phase solution gives $\Delta^* = K_{VH}e^{\mu_V} + V^{\rm tot}e^{-\mu_V}$. Simplifying with $H^{\rm tot*}$ and $\Delta^*$ gives

$$\frac{d}{dH^{\rm tot}}\frac{dH^{\rm tot}}{dt}\bigg|_{H^{\rm tot*}} = -\frac{e^{2\mu_V}K_{VH}^2}{(1 + K_{VH}e^{\mu_V})(K_{VH}e^{2\mu_V} + V^{\rm tot})}\ , \tag{25}$$

which is negative for all $K_{VH}$, $V^{\rm tot} > 0$ and $e^{\mu_V} < V^{\rm tot}$. Consequently, the fixed point is stable and training is guaranteed to converge.

### IV. UNSUPERVISED LEARNING IN MULTI-STABLE NETWORKS

The deterministic networks considered so far have a unique global fixed point, so their distribution over visible-species concentrations is a $\delta$-function regardless of hidden-species levels. This precludes unsupervised (generative) learning, whose goal is to tune the system's spontaneous distribution $P_H([V_1^{\rm tot}], \ldots, [V_n^{\rm tot}])$ to match a target distribution $Q([V_1^{\rm tot}], \ldots, [V_n^{\rm tot}])$ set by the environment, minimizing $D_{\rm KL}(Q\|P_H)$.

Multi-stable systems, however, naturally sample distributions over multiple attractor states, providing the expressivity needed for generative learning. During the wake phase, visible-species concentrations are fixed to samples from $Q$ using one of the targeted coupling mechanisms of Sec. II B; during the sleep phase, the system fluctuates freely. The autoregulatory learning rule then adjusts hidden-species levels to bring $P_H$ closer to $Q$. Below, we first develop the relevant ideas in a toy model, then consider a concrete example examined in the main text.

#### A. Toy 1D model

Consider a molecular system with dissipative dynamics (gene regulation, etc.) that make it multi-stable over several fixed points. Suppose it has a Lyapunov function $V(x)$ from which its dynamics are derived. Additionally, suppose that its dynamics are coupled to a training parameter, $\lambda$, whose contribution is derived from a potential. Consider its Lyapunov function

$$V(x, \lambda) = V_0(x) + \lambda x \tag{26}$$

With noise, the system naturally samples a distribution over the fixed points. For example, standard thermal noise $\eta$ can be used to generate Boltzmann statistics over the two fixed points:

$$\frac{dx}{dt} = \dot{V}(x, \lambda) + \eta \tag{27}$$

By adjusting $\lambda$ through training, we can change the system's distribution over fixed points $P_{\text{sys}}(x)$ to match a target one $P_{\text{target}}(x)$. After it is trained, the system will naturally act like a generative model, sampling new points from the learned distribution. Assuming the above dynamics, the system can be trained using the following training rule, where the training parameter is changed according to the average value of $x$ in a sleep and wake phase:

$$\frac{\partial \lambda}{\partial t} = -\frac{\partial V}{\partial \lambda}_{\text{sleep}} + \frac{\partial V}{\partial \lambda}_{\text{wake}} = \langle x \rangle_{\text{sleep}} - \langle x \rangle_{\text{wake}} \ . \tag{28}$$

#### B. Positive autoregulation example

Next, we consider a more concrete example – a basic formulation of bistable molecule $A$ under positive autoregulation:

$$\frac{d[A^{\text{mon}}]}{dt} = -\gamma [A^{\text{mon}}] + \frac{\alpha}{1 + (\frac{[A^{\text{mon}}]}{K})^{-n}} \ . \tag{29}$$

The two terms represent dynamics due to degradation and $n$-degree Hill production respectively, with degradation rate $\gamma$, production rate $\alpha$, and dissociation constant $K$. In the limit $n \to \infty$, so long as $\frac{\alpha}{\gamma} > K$, then the system hosts two stable fixed points at $[A^{\text{mon}}] = 0$ and $[A^{\text{mon}}] = \frac{\alpha}{\gamma}$. We are interested in tuning the relative time the system spends in its two fixed points by tuning $K$, which mechanistically could be achieved if $K$ is set by the fixed concentration of another molecule $C$.

Since we are in 1D, we can write $\frac{d[A^{\text{mon}}]}{dt} = f([A^{\text{mon}}]; K)$ and obtain $V([A^{\text{mon}}]; K) = \int_0^{[A^{\text{mon}}]} f(x; K)\ dx$ as the Lyapunov function for the dynamics. In order to evaluate $\frac{\partial V}{\partial K}$, we note that the only term that has $K$ dependence in $V$ is the integral of the cooperative production term. In the $n \to \infty$ limit, we can think of the production term as a step function $\alpha \Theta([A^{\text{mon}}] - K)$, in which case $\frac{\partial V}{\partial K} = -\alpha \Theta([A^{\text{mon}}] - K)$. Therefore, in this system, sleep-wake updates should follow the dynamics:

$$\frac{dK}{dt} \propto \langle \Theta([A^{\text{mon}}] - K) \rangle_{\text{sleep}} - \langle \Theta([A^{\text{mon}}] - K) \rangle_{\text{wake}} \ . \tag{30}$$

This has a simple interpretation - if the desired wake phase distribution is weighted more heavily towards the fixed point where $[A^{\text{mon}}] > K$, then one should decrease $K$, since this expands the basin of attraction around the $[A^{\text{mon}}] > K$ state. In order to tell whether or not the $[A^{\text{mon}}] > K$ fixed point is weighted more heavily in the wake phase distribution than in the sleep phase distribution, we can use the quantity $\Theta([A^{\text{mon}}] - K)$, since this serves as an indicator variable of the $[A^{\text{mon}}] > K$ fixed point. Therefore, if the wake phase distribution has more occurrences of the $[A^{\text{mon}}] > K$ than the sleep phase, then $\langle \Theta([A^{\text{mon}}] - K) \rangle_{\text{wake}} > \langle \Theta([A^{\text{mon}}] - K) \rangle_{\text{sleep}}$ and $\frac{dK}{dt} < 0$ as desired.

#### C. Gene toggle switch example

Finally, as a mechanistic picture of nonlinear positive autoregulation, we consider a simple gene toggle switch in which molecules $A$ and $B$ are transcription factors which repress each other's expression. The total concentration of molecule $A$ is described by the following dynamics:

$$\frac{d[A^{\text{tot}}]}{dt} = \frac{\alpha_A}{1 + (\frac{[B^{\text{mon}}]}{K_A})^n} - \gamma_A [A^{\text{tot}}] \ . \tag{31}$$

The dynamics of molecule $B$ are described by:

$$\frac{d[B^{\text{mon}}]}{dt} = \frac{\alpha_B}{1 + (\frac{[A^{\text{tot}}]}{K_B})^n} - \gamma_B[B^{\text{mon}}] - k_{\text{on}}[B^{\text{mon}}][C^{\text{mon}}] + k_{\text{off}}[BC] \ . \tag{32}$$

The concentration of $B$ is additionally affected by molecule $C$, which binds only to $B$ to form an inactive dimer, $BC$. The chemical kinetics are captured by the following dynamics:

$$\frac{d[C^{\text{mon}}]}{dt} = -k_{\text{on}}[B^{\text{mon}}][C^{\text{mon}}] + k_{\text{off}}[BC] \ , \tag{33}$$

$$\frac{d[BC]}{dt} = k_{\text{on}}[B^{\text{mon}}][C^{\text{mon}}] - k_{\text{off}}[BC] \ . \tag{34}$$

This system is bistable with two fixed points, which in the $n \to \infty$ limit consist of one where $[A^{\text{tot}}] = \frac{\alpha_A}{\gamma_A}$, $[B^{\text{mon}}] = 0$, $[C^{\text{mon}}] = [C^{\text{tot}}]$, $[BC] = 0$ and another where $[A^{\text{tot}}] = 0$, $[B^{\text{mon}}] = \frac{\alpha_B}{\gamma_B}$, $[C^{\text{mon}}] < [C^{\text{tot}}]$, $[BC] > 0$. The concentration of $C$ serves as a training parameter to modulate the probability distribution over the two states. For instance, when $C$ is present in large amounts, it will inactivate most $B$ molecules, leading the system to spend much more time in the $A$-dominant state.

To train this system to exhibit a desired fraction of time between its two fixed points, we follow the intuition suggested by our 1D model of positive autoregulation. There, we require a variable which can change the relative stability of the two fixed points, and another natural variable within the system that serves as an indicator variable for which fixed point the system is in. In this case, we know that increasing $[C^{\text{tot}}]$ increases the stability of the $[A^{\text{tot}}] > 0$ fixed point. By direct analogy to the 1D model Eq. 30, we can use $\Theta([B^{\text{mon}}] - K_A)$ as an indicator variable which is 1 when the system is in the $[A^{\text{tot}}] = 0$ fixed point. Following this logic, we arrive at a training rule:

$$\frac{d[C^{\text{tot}}]}{dt} \propto \langle \Theta([B^{\text{mon}}] - K_A) \rangle_{\text{sleep}} - \langle \Theta([B^{\text{mon}}] - K_A) \rangle_{\text{wake}} \ . \tag{35}$$

The above rule is analogous to the mathematically grounded rule given in the 1D setting, and is biologically plausible in the sense that the production of $A$ follows the same step functional form. However, we note that $[C^{\text{mon}}]$ serves equally well as an indicator of which fixed point the system is in; in the $[A^{\text{tot}}] > 0$ fixed point, $[C^{\text{mon}}] = [C^{\text{tot}}]$, while in the $[A^{\text{tot}}] = 0$ fixed point, $[C^{\text{mon}}] < [C^{\text{tot}}]$. Therefore, updates to $[C^{\text{tot}}]$ with the form

$$\frac{d[C^{\text{tot}}]}{dt} \propto \langle [C^{\text{mon}}] \rangle_{\text{wake}} - \langle [C^{\text{mon}}] \rangle_{\text{sleep}} \ . \tag{36}$$

will also have the same sign as Eq. 35 and hence will adjust the frequencies of the two fixed points within the system to the desired statistics. We therefore prefer to work with the Eq. 36 form of training rule, since it is both theoretically grounded but more similar to the training rules we derived for dimerization networks.

### V. HIGH-PASS FILTERING MECHANISMS ACROSS BIOLOGY

#### A. Temporal contrastive learning theory

In Sec. I we noted that the sleep–wake contrastive update

$$\frac{d\lambda_i}{dt} \propto \left\langle \frac{\partial E}{\partial \lambda_i} \right\rangle_{\text{sleep}} - \left\langle \frac{\partial E}{\partial \lambda_i} \right\rangle_{\text{wake}}$$

is not directly realizable in a molecular system unless one can store (and later take the difference between) sleep and wake gradients.

One way to approach the plausibility of Eq. 1 in molecular systems is to imagine that there is a switch between Hebbian and anti-Hebbian update mechanisms in coordination with the switch from wake to sleep phase. However, this proposal involves switching the sign of the training rule in conjuction with the sleep and wake environments, such an approach is difficult to coordinate autonomously on a cellular or molecular level.

We therefore use a temporal form of contrastive learning that avoids explicit memory storage or coordinated switching updates by replacing the difference of sleep-wake gradients with a *high-pass* filter:

$$\frac{d\lambda_i}{dt} \propto F_{HP}\left(\frac{\partial E}{\partial \lambda_i}\right) ,$$

as the environment toggles between sleep and wake. We can think of the high-pass filter as a mechanism that isolates the differences between the sleep and wake states, while ignoring their absolute values. Writing the synaptic signal as $s_i \equiv \langle \partial E/\partial \lambda_i \rangle$, the intuition behind the high-pass filter $F_{HP}$ is that it performs updates in proportion to a nonlinear function of the rate of change,

$$\frac{d\lambda_i}{dt} = g\left(\frac{ds_i}{dt}\right),$$

with $g$ monotone and threshold-like so that slow variations contribute weakly. We assume the sleep-to-wake transition is fast while wake-to-sleep relaxation is slow; then filtering emphasizes the fast jump in $s_i$ while suppressing the slow return, effectively isolating $s_i^{\text{sleep}} - s_i^{\text{wake}}$.

More concretely, we can think of $F_{HP}$ as being the result of a system which exhibits integral feedback control. Let $\lambda_i$ be produced/removed by a molecular species $u_i$, and let $\frac{d\lambda_i}{dt} = g(u_i)$, where $u_i$ is coupled to $s_i$ and placed under integral feedback. A standard reduced model [6] is

$$\tau_u \frac{du}{dt} = -u + k\,(s(t) - m), \tag{37}$$

$$\tau_m \frac{dm}{dt} = (u - u_0), \tag{38}$$

which implies the driven overdamped oscillator form

$$\tau_u \frac{d^2u}{dt^2} = -\frac{du}{dt} + k\left(\frac{ds}{dt} - \frac{1}{\tau_m}(u - u_0)\right), \tag{39}$$

with $u$ driven by $\dot{s}(t)$. For a linear ramp $s(t) = s\,t$ (and $u_0 = 0$), the steady-state response is $u = \tau_m s$.

Now assume a stereotyped forcing where $s(t)$ changes over a fast timescale $\tau_f$ and relaxes over a slow timescale $\tau_s \gg \tau_f$, as in bursty influx followed by slow decay. During the fast transition,

$$s(t) = s^{\text{sleep}} + \frac{t}{\tau_f}\left(s^{\text{wake}} - s^{\text{sleep}}\right), \qquad u(t) \approx \frac{s^{\text{wake}} - s^{\text{sleep}}}{\tau_f},$$

so if $d\lambda/dt \propto u$ then over the fast window

$$\int_0^{\tau_f} u\,dt = s^{\text{wake}} - s^{\text{sleep}}.$$

Over a full cycle, the slow relaxation would cancel this contribution. However, if the update is gated/weighted toward large $u$ (equivalently, via thresholding nonlinearity $g$), then in the limit $\tau_f \ll \tau_s$ one obtains [2]

$$\int_0^{\tau_f + \tau_s} g(u)\,dt \propto s^{\text{sleep}} - s^{\text{wake}}. \tag{40}$$

Thus, sleep–wake contrastive learning can be implemented in molecular systems without explicit memory storage, using integral feedback combined with natural temporal asymmetry in environmental fluctuations.

#### B. Molecular realizations

Our proposed training rule requires a regulatory response to rapid changes in species concentrations but not to slow changes in those same concentrations (high-pass filtering). This functional capability of filtering out steady-state backgrounds while responding to fluctuations is a common feature of cellular decision-making. It is a property of diverse circuit motifs ranging from integral feedback to physical processes like phase separation. Here we categorize these mechanisms by their underlying control-theoretic or physical principle rather than by organism, illustrating the ubiquity of the rate-sensitivity required for our proposed training rule.

##### *1. Integral Feedback Control*

One of the most stringent ways to implement a high-pass filter in biology is through integral feedback. In this motif, a specific molecular species acts as a "memory" variable that accumulates (integrates) the error between the system's current state and its set point. As long as an error exists, the memory variable continues to change, driving a negative feedback loop that eventually forces the error to zero. This mathematical structure ensures that the system perfectly adapts to constant inputs (zero steady-state output) while responding sensitively to rapid changes, i.e., the response is set by the time derivative of the input. Such a time-derivative operation is naturally a high-pass filter [7].

- Bacterial Chemotaxis (CheR/CheB System): This is a canonical example of biological integral control. The activity of the chemoreceptor complex controls the kinase CheA. However, the receptor's activity also drives a slow methylation modification cycle mediated by the methyltransferase CheR and the methylesterase CheB. The methylation level of the receptor effectively integrates the difference between ligand occupancy and the system's activity target. This allows *E. coli* to sense chemical gradients (changes in concentration) while adapting to background concentrations spanning several orders of magnitude [6, 8, 9].
- Yeast Osmoregulation (HOG Pathway): In *S. cerevisiae* and other fungi [10], the High Osmolarity Glycerol (HOG) pathway maintains turgor pressure. Upon osmotic shock, the Hog1 kinase activates, driving the production and retention of glycerol. Intracellular glycerol accumulates until turgor pressure is restored, at which point the mechanical stress sensor turns off Hog1. Here, the intracellular glycerol concentration serves as the physical integration variable, accumulating until it cancels out the environmental osmotic stress [11, 12].
- Calcium Homeostasis: In mammalian cells, cytosolic calcium spikes trigger immediate signaling events but also activate high-affinity pumps (such as SERCA and PMCA) that sequester calcium into the ER or eject it from the cell. These pumps operate to restore calcium to a low homeostatic set point. The total amount of calcium pumped matches the integral of the influx, effectively filtering out sustained increases in flux while allowing transient signaling spikes to propagate [13].

##### 2. *Incoherent Feedforward Loops*

The Incoherent Feedforward Loop (IFFL) is a network motif where an input signal splits into two paths: a direct activation path and an indirect inhibition path that acts on the same output node. Crucially, if the inhibition path is delayed relative to the activation path, a step increase in input triggers a transient pulse of output. The activator arrives first, followed later by the inhibitor which shuts the response down. This motif is a standard mechanism for "Fold-Change Detection", where the system's peak response depends on the relative rate of change rather than the absolute concentration [14].

- NF-$\kappa$B Signaling: In the immune response to cytokines (e.g., TNF$\alpha$), the activation of the transcription factor NF-$\kappa$B leads to the transcription of its own inhibitor, I$\kappa$B$\alpha$. This delayed negative feedback loop (which functions similarly to an IFFL when viewing the cytokine as the input) causes NF-$\kappa$B activity to pulse in response to step inputs. The system effectively takes a time-derivative of the cytokine concentration, filtering out slowly increasing levels [15–17].
- MicroRNA-Mediated Buffering: Many microRNAs target the same transcripts that drive their own expression, or are co-expressed with their targets in IFFL topologies. This allows the system to dampen steady-state protein levels (buffering noise) while allowing rapid transitions in gene expression to propagate, effectively functioning as a high-pass filter for gene regulation [18–20].

##### 3. *Receptor Desensitization (Conformational Switching)*

Adaptation can occur at the level of single macromolecules through thermodynamic conformational dynamics. In this mechanism, a receptor possesses three states: resting, active, and desensitized (inactive but high-affinity). If the desensitized state is thermodynamically favored under conditions of prolonged ligand binding, the population of signaling-active receptors will strictly be transient. The system generates a signal only during the non-equilibrium shift from resting to desensitized, effectively filtering out sustained ligand presence.

- Ligand-Gated Ion Channels: Neurotransmitter receptors, such as Nicotinic Acetylcholine Receptors (nAChRs) and GABA$_A$ receptors, open rapidly upon ligand binding but spontaneously transition to a closed, desensitized conformation if the ligand remains bound for milliseconds to seconds [21]. This ensures that the postsynaptic response is triggered only by the arrival of the signal and not prolonged stimulation.

- G-Protein Coupled Receptors (GPCRs): Upon activation by a ligand, GPCRs are rapidly phosphorylated by G-protein coupled receptor kinases (GRKs) and subsequently bound by $\beta$-arrestins [22, 23]. This binding uncouples the receptor from G-proteins (desensitization) and triggers internalization. This molecular process serves as a high-pass filter for responses to changes in hormone concentration.

##### *4. Phase Separation*

Recent work [24] has highlighted that passive physical phase transitions can act as dynamic filters without complex regulatory wiring. Biomolecular condensates formed by liquid-liquid phase separation (LLPS) coexist with a dilute phase. The dense phase acts as a reservoir that exchanges molecules with the dilute phase at a finite rate. This exchange buffering effectively "absorbs" low-frequency fluctuations, while high-frequency fluctuations occur too rapidly to equilibrate and are thus expressed in the dilute phase concentration.

### VI. TIMESCALES FOR LEARNING AND LONG-TERM MEMORY

#### A. Timescales in molecular sleep-wake

The learning framework introduced here involves multiple timescales – a slow timescale of clamping relaxation $\tau_s$, a fast timescale of clamping application $\tau_f$, a timescale associated with the memory kernel $\tau_K$, as well as an implicit timescale of system equilibration, $\tau_{eq}$. A sufficient condition for temporal contrastive learning to work is that $u$ defined by Eq. 39 should approximate the time-derivative of the (equilibrium) synaptic signal $s_{ij}$. This implies $\tau_K < \tau_s, \tau_f$. Performance of temporal contrastive learning is further improved when the nonlinearity $g$ in Eq. 40 can easily separate the convolved signal $u_{ij}$ during the slow and fast ramps of the timescale. This separation of the two ramps is therefore achieved more easily when $\tau_f < \tau_s$. Finally, the assumption of equilibrium synaptic signal being fed into the kernel $K$ implies that $\tau_{eq} < \tau_K$. Putting these relations together, we arrive at the following sufficient regime for good performance of temporal contrastive learning:

$$\tau_{eq} < \tau_K < \tau_f < \tau_s \ . \tag{41}$$

It is possible to relax the constraints on $\tau_f > \tau_K$. In this case, viewed on the timescale of $\tau_K$, the fast ramp appears as a step function, and again Eq. 40 holds.

#### B. Long-term memory of learned behavior

Although the tunability of hidden species concentrations makes them effective training parameters, it also creates a tension: the same malleability that enables learning also makes learned states vulnerable to erasure. Degradation, stochastic fluctuations, and dilution through cell division can all erode hidden species levels over time. The severity of this problem depends strongly on context. We therefore discuss strategies for maintaining learned behavior separately for systems with minimal dilution (e.g., *in vitro* circuits or non-dividing cells) and for rapidly dividing cells where concentrations are halved at each division.

**Non-dividing or slowly dividing systems:** When dilution is not a concern, learned concentrations can be maintained through approaches already explored in experiments. Here, a primary concern is a baseline level of protein degradation common to many molecular realizations of integral feedback control.

*Separation of training and testing phases.* If the learning circuit requires continual protein degradation, then the simplest approach to avoid issues from protein degradation is to turn off the learning circuit once the desired behavior has been acquired. If autoregulatory feedback is deactivated after training, learned hidden species levels will persist as long as other endogenous sources of degradation or fluctuation are slow compared to the timescale on which the behavior needs to be maintained. One could also transfer the information stored in hidden species concentrations into a chemically stable set of molecules not subject to autoregulatory dynamics, further protecting the memory from perturbation. This is directly analogous to freezing weights after training a neural network, and is especially natural for engineered in vitro systems such as DNA-based molecular circuits [25].

*Continual learning with slow forgetting.* Alternatively, one can leave the learning circuit active but ensure that the learning timescale is long by slowing both production and degradation timescales for the hidden species. The system can then be "tested" on timescales shorter than this learning timescale without appreciably forgetting what it has learned. Such slow learning (due to biochemically long-lived hidden species) is acceptable when environmental statistics are stable. Such a system can also gradually adapt if environmental correlations shift, without requiring an explicit retraining phase.

**Rapidly dividing cells:** Cell division poses a qualitatively different challenge because each division roughly halves hidden species concentrations. Without a restorative mechanism, learned states will be diluted away within a few generations.

*Continual reinforcement from persistent environmental correlations.* If the environmental statistics that drove training continue to be present during subsequent divisions, the autoregulatory learning rule will continuously push hidden species levels back toward their learned values, counteracting dilution. Each generation of cells effectively re-learns the same associations from ongoing environmental input. This approach does require that the generation time is not so fast that the autoregulatory feedback cannot keep up with the rate of dilution.

*Pre-programmed attractors for hidden species.* A more robust solution is to couple the learning dynamics to a pre-existing multistable circuit that defines discrete attractor states for hidden species concentrations. Evolution could program a set of allowed expression patterns, corresponding to stable attractors, as is the case with genes that specify cell type. The learning rule would then select whichever attractor produces input-output behavior closest to what the environment demands. Once the system settles into the appropriate attractor, the multistable circuit actively restores hidden species levels against dilution, as long as the perturbation from division is not large enough to kick the system out of its attractor basin. A tradeoff in this approach is that hidden species levels are effectively discretized, and hence more hidden species will be required to get the same level of expressivity reported in this paper.

*Epigenetic memory.* Perhaps the most biologically natural route to long-term memory in dividing cells is through epigenetic modifications: heritable chemical marks on DNA and histones that regulate gene expression without altering the underlying genetic sequence. Modifications such as DNA methylation and repressive histone marks (H3K9me3, H3K27me3) are actively maintained through cell division by dedicated enzymatic machinery (e.g., DNMT1 for maintenance methylation) and can stably propagate gene expression states across many generations [26]. If hidden species expression is controlled by epigenetically regulated loci, then learned concentrations could be encoded as stable epigenetic states that persist through division.

Conventionally, epigenetic memory has been understood as binary: genes are locked into either a fully silenced or fully active state through self-reinforcing feedback between DNA methylation and histone modifications. Under this view, the number of stably maintainable hidden species levels would be limited to a discrete set (on/off), which would limit the expressivity of learned behaviors. But this limitation can be alleviated by increasing the number of hidden species.

An alternative solution is suggested by recent work on the analog [27] nature of epigenetic memory in some conditions, i.e., cells can stably maintain gene expression at a continuous range of intermediate levels, not just the two extremes. The mechanism involves graded DNA methylation: the mean fraction of methylated CpGs at a gene's promoter can be set to intermediate values that remain stable across cell divisions and that correlate inversely with expression level. If the hidden species levels learned through autoregulatory learning dynamics can be written into graded methylation states at promoters controlling hidden species production, then transient concentration-based memory could be converted into a durable epigenetic record that survives through cell divisions.

### VII. COMPUTATIONAL DETAILS

#### A. Simulating networks

##### 1. *Modeling dimerization*

We simulate our training protocol in a minimal model of a protein dimerization network, the simplest biologically relevant example of a many-to-many molecular network. Dimerization networks consist of $N$ different species of monomers that form homodimers and heterodimers with all other monomer species present. The formation of larger complexes, such as trimers, is not considered.

$$m_i + m_j \rightleftharpoons d_{ij}$$

All reactions are stoichiometric, and the extent of dimerization between each monomer pair in equilibrium is determined by mass action kinetics and encoded in an equilibrium binding constant $K_{ij}$.

$$K_{ij} = \frac{[d_{ij}]}{[m_i][m_j]}$$

#### 2. Parameters

Each network we simulated was fully connected, with random concentrations and binding constants sampled from log-normal distributions. Concentrations were initialized with values ranging from $10^{-3} - 10^{3}$ and binding constants ranging from $10^{-5} - 10^{7}$. Both quantities are dimensionless because we normalize concentrations by the number density of the solvent (assumed to be water at standard temperature and pressure). These parameter ranges were adopted from previous work on computation through dimerization [28], which discusses their consistency with the natural range of concentrations and binding affinities observed in biology.

#### 3. Packages

When training networks for Pavlovian conditioning and supervised learning, we computed the input-output functions of networks using the Python Equilibrium Toolkit (EQTK) package, which efficiently solves for the equilibrium concentrations of biochemical reaction networks through energy minimization, which is defined as the solution to

$$\begin{cases} m_i^{\text{tot}} & = m_i + 2d_{ii} + \sum_{j\neq i} d_{ij} \\ K_{ij} & = \frac{d_{ij}}{m_i m_j} \end{cases} \tag{42}$$

We consider the function computed by the network to be the response of the output molecule concentration to changes in the total input molecule concentration. To determine this response function, input molecules were titrated over concentrations ranging from $10^{-3} - 10^{3}$, with fixed concentrations of hidden molecules, and EQTK was used to compute the corresponding equilibrium concentration of the output molecule at each titration point.

### B. Training details

#### 1. Training protocol

As described in Section I, our practical training protocol alternates between two phases: a sleep phase, where the environment sets concentrations of a subset of visible species, designated as inputs, and a wake phase, where the environment sets concentrations of all input and output visible species.

*a. Sleep phase* After initializing networks and selecting training data, we began training with the sleep phase. In this phase, we used EQTK to find equilibrium concentrations while fixing total concentrations of input, hidden, and output species. This is equivalent to solving

$$\begin{cases} m_i^{\text{tot}} & = m_i + 2d_{ii} + \sum_{j\neq i} d_{ij} \quad \forall i \\ K_{ij} & = \frac{d_{ij}}{m_i m_j} \end{cases} \tag{43}$$

*b. Wake phase* Next, we ran the wake phase. In this phase, input and hidden species retain the same total concentrations as in the sleep phase, but total concentrations of output species are adjusted to a new value to set their monomer concentrations to the training target $m_0^*$. The set of equations corresponding to the wake phase is

$$\begin{cases} m_i^{\text{tot}} & = m_i + 2d_{ii} + \sum_{j\neq i} d_{ij} \quad \forall i > 0 \\ m_0 & = m_0^* \\ K_{ij} & = \frac{d_{ij}}{m_i m_j} \end{cases} \tag{44}$$

*c. Training update* After computing sleep and wake equilibria, we updated hidden species concentrations by an amount equal to the difference between their sleep and wake monomer levels according to our proposed learning dynamics,

$$\Delta[m_i^{\text{tot}}] = \eta\big(\langle \ln[m_i^{\text{mon}}]\rangle_{\text{wake}} - \langle \ln[m_i^{\text{mon}}]\rangle_{\text{sleep}}\big), \tag{45}$$

where $\eta$ is a fixed learning rate. Updates were made to sleep phase concentrations, which were assumed to be the baseline concentrations that the network returns to after experiencing a temporary concentration shift in the wake phase. When training data consisted of multiple points, the sleep and wake phases were repeated independently for each training point and Eq. 45 was averaged to obtain a single update per training epoch. We repeated the sleep-wake-update cycle until hidden species concentrations converged. Input and output visible species concentrations were not updated and remained fixed at their initial values throughout training.

#### 2. *Clamping concentrations*

Inputs were chosen to be total concentrations, which we fixed by imposing a mass constraint on the input visible species. In contrast, we generally chose outputs to be monomer concentrations of visible species. In practice, to fix these concentrations we removed the columns corresponding to the output visible species from the network's stoichiometry matrix, along with the rows corresponding to their homodimerization reactions. Then, we replaced all equilibrium constants $K_{0j}$ for reactions that involved the output visible species with effective equilibrium constants,

$$\tilde{K}_{0j} = K_{0j} \cdot [m_0^*].$$

In doing so, we assumed that the dimerization network was in contact with a reservoir of the visible output species $m_0$ which buffers its concentration to $[m_0^*]$, while respecting the equilibrium constraints of the other species. This amounts to assuming that the system is open to the output species, so that its total concentration is no longer conserved and it is effectively in a grand-canonical ensemble, while all other species remain subject to mass constraints.

### C. Choosing training targets

Dimerization networks are not expressive enough to represent arbitrary input-output functions. The parameters of a network, namely its size and binding constants, restrict the class of functions it can implement [28]. As a result, meaningful training targets must be chosen from the accessible space of functions for a given network.

To identify such targets, we searched over trainable parameters by varying the concentrations of hidden species while keeping network topology and binding constants fixed. Specifically, we sampled total concentrations of hidden species from a log-uniform distribution, ranging from $10^{-3} - 10^{3}$, and plotted the output function corresponding to each parameter set. From these candidates, we chose two sets of parameters that led to qualitatively different behavior, such as monotonicity (Fig. 3b) or logic structure (Fig. 3c).

We then used one of these functions as a training target and applied our learning dynamics to adjust concentrations of the other so that the network matched the target output. We selected training data manually by choosing 1-5 points from the target function that captured its defining features.

### D. Figure descriptions

#### 1. *Pavlovian conditioning*

In Fig. 2b, we performed Pavlovian conditioning of a 3-species dimerization network following the framework described in Sec. II A 1. The network consists of two visible species $V_B$ (bell) and $V_F$ (food) and one hidden species $H$, coupled by fixed pairwise binding affinities $K_{ij}$. We initialized our network with species concentrations $[H^{\text{tot}}] = 455.34$ and $[V_F^{\text{tot}}] = 50.0$. In both sleep and wake phases, the bell monomer level was fixed at $[V_B^{\text{mon}}] = 6.51 \times 10^{-2}$. Initially, this resulted in a network which output $[V_F^{\text{mon}}] = 1.931 \times 10^{-2}$, but the target response was $[V_F^{\text{mon}}] = 1.169$. Therefore, we trained our our network for $11,500$ iterations with a learning rate $\eta = 0.02$. Interaction constants $K_{ij}$ were sampled from a log-uniform distribution, ranging from $10^{-7} - 10^{5}$. Following training, we arrived at a network with species concentrations $[H^{\text{tot}}] = 51.09$, which outputs $[V_F^{\text{mon}}] = 1.168$.

#### 2. *Step-down to step-up*

In Fig. 3a, we performed supervised training of a 3-species dimerization network following the framework described in Sec. III. We initialized our step-down network with total species concentration values and $K_{ij}$ taken from the example shown in Fig. 3b of [28]. We then train our network for $10,000$ iterations with a learning rate of $\eta = 1$. We chose our target values by computing the equilibrium concentrations of $[V_F^{\text{mon}}]$ for $[V_B^{\text{tot}}] = 0.01$ and 10 for the step-up function in [28].

#### 3. *Monotonic to non-monotonic*

In Fig. 3b, we performed supervised training of an 8-species dimerization network following the framework described in Sec. III. We initialized our network with total species concentration values chosen from a log-uniform distribution ranging from $10^{-3}$ to $10^{3}$, and $K_{ij}$ chosen from a log-uniform distribution ranging from $10^{-5}$ to $10^{7}$. We then trained our network for 50,000 iterations with a learning rate of $\eta = 90$. We chose our target function by generating 25 sets of randomly chosen hidden species concentrations and computing input-output curves for each one. One target output was selected, and three training points were selected from the curve, one at a low input concentration, one at a medium input concentration (corresponding to the maximum of the non-monotonic target function) and one at a high input concentration.

##### *4. Higher dimensional training*

In Fig. 3c, we performed supervised training of a 6-species dimerization network following the framework described in Sec. III. Here, we selected $V_1^{\text{tot}}$ and $V_2^{\text{tot}}$ as inputs to the molecular network. We initialized our network with total species concentration values chosen from a log-uniform distribution ranging from $10^{-3}$ to $10^{3}$, and $K_{ij}$ chosen from a log-uniform distribution ranging from $10^{-5}$ to $10^{7}$. We then trained our network for $20,000$ iterations with a learning rate of $\eta = 0.1$. To choose our target function, we randomly generated 25 sets of hidden species concentrations using Latin Hypercube Sampling and plotted their outputs, choosing as a target function one that was qualitatively different from the initial one. Five training points were selected from the target function, four corresponding to combinations of high and low input concentrations, and one at an intermediate value of both inputs.

##### *5. Iris classification*

In Fig. 3d, we used a 12-species dimerization network to classify the Iris dataset. The Iris dataset is composed of 150 data points, evenly divided among three different species of irises – *setosa*, *versicolor*, and *virginica*. Each data point consists of four features – petal length, petal width, sepal length, and sepal width, which characterize one of the three species. We restricted our analysis to a binary classification task using the 100 samples that belong to the *setosa* and *versicolor* classes.

We selected molecular species $V_1$, $V_2$, $V_3$, and $V_4$ as input species and species $V_5$ as output species. We encoded Iris inputs into the molecular network by setting the concentrations of the input molecular species to $10^{x}$, where $x$ was the feature value. The network output was taken to be the monomer concentration of $V_5$. Outputs above the threshold of 0.1 were classified as *setosa*, while outputs below the threshold were classified as *versicolor*.

We initialized the network with total species concentrations and $K_{ij}$ chosen from log-uniform distributions ranging from $10^{-3}$ to $10^{3}$ and $10^{-5}$ to $10^{7}$, respectively. We then trained our network for $10,000$ iterations with a learning rate of $\eta = 0.1$. We randomly selected 80% of the data to train on and reserved the remaining 20% for testing.

We tracked classification accuracy, defined as the fraction of data points correctly classified based on whether the output monomer concentration exceeded the 0.1 threshold, for both training and test data throughout training.

##### *6. Generative learning*

In Fig. 4c, we simulated a bistable gene toggle switch with two mutually-inhibiting species $A$ and $B$. We introduced a third species, $H$, which serves as a training parameter. Species $H$ reversibly binds with $B$ to form an inactive complex $BH$, thereby modulating the relative occupancy of the two fixed points of the system.

Species $A$ and $B$ are produced at rates $\alpha_A = \alpha_B = 12.5$. Each represses the other's production through a Hill function with exponent $n_A = n_B = 4$ and dissociation constants $K_A = K_B = 1.5$. Both species undergo linear degradation at rates $\gamma_A = \gamma_B = 2$. There is mass conservation of $H^{\text{tot}}$, where $[H^{\text{tot}}] = [H^{\text{mon}}] + [BH]$. At steady state, we assumed a quasi-equilibrium between $B$ and $H$, with binding constant

$$K_{\text{BH}} = \frac{[BH]}{[B^{\text{mon}}][H^{\text{mon}}]} = \frac{k_{\text{on}}}{k_{\text{off}}} = \frac{2}{2} = 1.$$

To induce stochastic switching between the two stable states, we multiplied each rate $k \in \{\gamma_A,\ \gamma_B,\ \alpha_A,\ \alpha_B\}$ by a log-normal noise term $\eta(t)$ at each timestep,

$$k_{\text{noisy}}(t) = k \cdot \eta(t), \quad \eta(t) \sim \text{LogNormal}(\mu = 0, \sigma = \sigma_{\text{noise}}).$$

Here, $k_{\text{noisy}}$ is the stochastically perturbed version of the baseline rate $k$. For each stochastic simulation, we set $\sigma_{\text{noise}} = 1.2$ to observe frequent switching between the bistable states within a single run.

We determined an appropriate simulation length by running stochastic simulations ($dt = 0.01$ and $H^{\text{tot}} = 0$) with increasing numbers of timesteps until the steady state distributions converged. Convergence was assessed by requiring all pairwise Wasserstein distances among 10 independent runs to fall within ∼1-10% of the support width. Based on this analysis, we ran each stochastic simulation for $10^7$ timesteps.

Our training data was drawn from a distribution $P^{\text{env}}([A^{\text{tot}}])$. We generated $P^{\text{env}}([A^{\text{tot}}])$ by running a stochastic simulation with the parameters described above and $[H^{\text{tot}}] = 4.0$.

We initialized the system with $[A^{\text{tot}}] = [B^{\text{mon}}] = 3.0$, $[BH] = 0$ and $[H^{\text{tot}}] = 0.1$. During each sleep phase simulation, we ran stochastic dynamics with $dt = 0.01$ for $10^7$ timesteps and computed $\langle[H^{\text{mon}}]\rangle$ as the time-averaged concentration of unbound $H$.

In each wake phase simulation, we independently drew 100 samples of $([A^{\text{tot}}], [B^{\text{tot}}])$ from the target distribution $P^{\text{env}}([A^{\text{tot}}])$. For each sample, we computed $[H^{\text{mon}}]$ at the current level of $[H^{\text{tot}}]$ by assuming quasi-equilibrium between $B$ and $H$ and running deterministic dynamics ($\sigma_{\text{noise}} = 0$) for 1000 timesteps with $dt = 0.01$. We computed $\langle[H^{\text{mon}}]\rangle_{\text{wake}}$ as the average unbound $H$ across these samples. Then, we updated $[H^{\text{tot}}]$ based on the difference between sleep and wake averages.

We trained the system for 1,000 epochs with a learning rate of $\eta = 0.25$. In the end, the trained system converged to $[H^{\text{tot}}] = 2.74$.

#### 7. *Trimerization networks*

In Fig. 5a, we performed supervised training of 10-species fully-connected dimerization and trimerization networks following the framework described in Sec. III. We initialized the dimerization network with concentrations and $K_{ij}$ sampled from log-uniform distributions ranging from $10^{-3} - 10^3$ and $10^{-5} - 10^7$, respectively. We then generated a trimerization network that was a copy of the dimerization network with several added trimerization reactions, specifically self-trimerization of species $V_2$ and $H_5$, and hetero-trimerization between species $V_1$, $V_2$, and $H_5$. We sampled trimerization equilibrium constants $K_{ijk}$ from a log-uniform distribution ranging from $10^{-5} - 10^7$ and assumed that trimers form directly from monomers, without forming dimer intermediates.

We trained both networks for 70,000 iterations with a learning rate of $\eta = 10$ on identical training data, chosen by sweeping over hidden species concentrations in the dimerization network and selecting representative input-output pairs.

#### 8. *Networks with unmodeled interactions*

In Fig. 5b, we performed supervised training of a fully-connected 5-species dimerization networks following the framework described in Sec. III. We initialized our network with total species concentrations and $K_{ij}$ sampled from log-uniform distributions ranging from $10^{-3} - 10^3$ and $10^{-5} - 10^7$, respectively. We designated two of the species as input and output visible species, one as a hidden species, and the remaining two as species that have unexpected interactions with the network and whose concentrations do not get trained by the training rule. We then trained our network for 200,000 iterations with a learning rate $\eta = 0.9$. We chose our training data by sweeping over hidden species concentrations and selecting representative input-output pairs.

#### 9. *Learning in heterogeneous populations*

In Fig. 5c, we performed supervised training of an ensemble of 40 fully-connected 6-species dimerization networks following the protocol described in Sec. III. We initialized each network with the same total concentrations, sampled from a log-uniform distribution ranging from $10^{-3} - 10^3$, but different $K_{ij}$, all sampled from a log-uniform distribution ranging from $10^{-5} - 10^7$. We trained each network for 8,000 iterations with a learning rate of $\eta = 0.8$. We used $V_1^{\text{in}} = 0.1$, $V_2^{\text{out}} = 1$ as the training target for each network.

#### 10. *Alternate clamping rules*

In Fig. 6a(i), we performed supervised training with nudging of a fully-connected 5-species dimerization network. We initialized the total concentrations of each species and $K_{ij}$ by sampling log-uniform distributions ranging from $10^{-3} - 10^3$ and $10^{-5} - 10^7$, respectively. While the sleep phase of training proceeded as usual (as described in Sec. III), in the wake phase, we did not clamp the monomer concentration of $V_2^{\text{out}}$ to the exact desired response but instead nudged it in the desired direction. To do so, we first determined the current monomer concentration with EQTK. Then we computed a directional term by comparing it to the target output – a negative sign if the current output was too high and a positive sign if the current output was too low. Using this direction, we shifted the value of the output species up or down by a small, proportional amount by adding or subtracting a fraction $c$ of the current output in the wake phase. In this way, we adaptively "nudged" the system towards the target rather than fixing it there at each iteration of the wake phase. We trained with four different nudging strengths, using $c = 0.01, 0.1, 0.5$, and 0.8. We ran each trial for 80,000 iterations with a learning rate of $\eta = 0.1$. In each training run, we chose $V_1^{\text{in}} = 0.1,\ V_2^{\text{out}} = 1$ as our target.

##### *11. Alternate update dynamics*

In Fig. 6a(ii), we performed supervised training of a fully-connected 10-species dimerization network. We initialized the total concentrations of each species and $K_{ij}$ by sampling log-uniform distributions ranging from $10^{-3} - 10^{3}$ and $10^{-5} - 10^{7}$, respectively. Training proceeded as described in III, except that hidden species concentrations were updated after each sleep and each wake phase, rather than based on the difference between sleep and wake phase hidden monomer concentrations. Specifically, hidden species concentrations were decreased following each sleep phase and increased following each wake phase, in proportion to the corresponding hidden species monomer concentrations. We trained the same network with combined and separate sleep-wake updates. We trained networks for 50,000 iterations with a learning rate of $\eta = 0.1$. We chose $V_1^{\text{in}} = 0.01,\ V_2^{\text{out}} = 1$ as our training data for both runs.

##### *12. Alternate training rules*

In Fig. 6b, we performed supervised training of a 10-species dimerization network according to the framework described in Sec. III. We initialized our network with total concentrations and $K_{ij}$ sampled from log-uniform distributions ranging from $10^{-3} - 10^{3}$ and $10^{-5} - 10^{7}$, respectively. We trained the network using three different training rules: a log update,

$$\Delta H_i^{\text{tot}} \propto \ln\langle H_i^{\text{mon}}\rangle_{\text{sleep}} - \ln\langle H_i^{\text{mon}}\rangle_{\text{wake}},$$

a linear update,

$$\Delta H_i^{\text{tot}} \propto \langle H_i^{\text{mon}}\rangle_{\text{sleep}} - \langle H_i^{\text{mon}}\rangle_{\text{wake}},$$

and a Hill function update,

$$\Delta H_i^{\text{tot}} \propto \left\langle \frac{(H_i^{\text{mon}})^2}{1 + (H_i^{\text{mon}})^2} \right\rangle_{\text{sleep}} - \left\langle \frac{(H_i^{\text{mon}})^2}{1 + (H_i^{\text{mon}})^2} \right\rangle_{\text{wake}}.$$

We trained each network for 20,000 iterations with a learning rate $\eta = 0.5$. We selected $V_1^{\text{in}} = 0.01,\ V_2^{\text{out}} = 1$ as the training target for each network.

[1] D. H. Ackley, G. E. Hinton, and T. J. Sejnowski, Cognitive science **9**, 147 (1985).
[2] M. J. Falk, A. T. Strupp, B. Scellier, and A. Murugan, Nature Communications **16**, 2163 (2025).
[3] W. Poole, A. Ortiz-Munoz, A. Behera, N. S. Jones, T. E. Ouldridge, E. Winfree, and M. Gopalkrishnan, in *International conference on DNA-based computers* (Springer, 2017) pp. 210–231.
[4] W. Poole, T. E. Ouldridge, and M. Gopalkrishnan, Journal of the Royal Society Interface **22**, 20240373 (2025).
[5] B. Scellier and Y. Bengio, Frontiers in computational neuroscience **11**, 24 (2017).
[6] T.-M. Yi, Y. Huang, M. I. Simon, and J. Doyle, Proceedings of the National Academy of Sciences **97**, 4649 (2000).
[7] K. J. Åström and R. Murray, *Feedback systems: an introduction for scientists and engineers* (Princeton university press, 2021).
[8] N. Barkai and S. Leibler, Nature **387**, 913 (1997).
[9] U. Alon, M. G. Surette, N. Barkai, and S. Leibler, Nature **397**, 168 (1999).
[10] T. You, P. Ingram, M. D. Jacobsen, E. Cook, A. McDonagh, T. Thorne, M. D. Lenardon, A. P. De Moura, M. C. Romano, M. Thiel, *et al.*, BMC research notes **5**, 258 (2012).
[11] D. Muzzey, C. A. Gómez-Uribe, J. T. Mettetal, and A. van Oudenaarden, Cell **138**, 160 (2009).
